# Mitochondria Exhibit Changes in Morphology/Function to Support Increased Glutamate Release in Tau_P301L_ Neurons Prior to Reduction in Presynaptic Vesicle Release

**DOI:** 10.1101/2023.06.30.547132

**Authors:** Rachel Cotter, Morgan Hellums, Delaney Gray, David Batista, Jeremiah Pfitzer, Miranda N. Reed, Michael W. Gramlich

## Abstract

We have shown that tauopathy models display early-stage hyperexcitability due to increased presynaptic glutamate release that is mediated by an increase in vesicular glutamate transporter-1 (VGlut1). This hyperexcitability increases energy demand which in turn would increase demand on mitochondria. It is unclear, however, how early-stage presynaptic changes in glutamate release are supported by or influence the function of mitochondria. Using Large Area Scanning Electron Microscopy (LA-SEM) and fluorescence microscopy, we demonstrate that mitochondrial changes in morphology, structure, and function in CA1/CA3 hippocampal neurons decrease resting mitochondrial membrane potential in P301L mice. However, P301L mitochondria maintain a high membrane potential during levels of high activity, suggesting that they can support increased energy demand during hyperexcitability. These activity-dependent differences in membrane potential can be rescued by inhibiting ATP-dependent VGlut1 vesicle refilling. This indicates that the increased VGlut1 per vesicle observed in P301L mice contributes to the differences in mitochondria membrane potential. Notably, the mitochondrial dysfunction in P301L mice occurs before any observable alterations in presynaptic release mechanics, suggesting these changes may represent early therapeutic targets. Finally, we propose a model of increased glutamate-mediated changes in mitochondrial morphology and function in P301L neurons that represents a potentially targetable pathway to reduce or arrest neurodegeneration.

## Introduction

Tauopathies are a class of conditions that result from hyperphosphorylation and aggregation of tau that leads to neurodegeneration, such as Alzheimer’s disease (AD), Parkinson’s disease (PD), Pick’s disease, and Frontotemporal dementia with Parkinsonson-17 (FTDP-17). Synaptic communication decreases caused by reduced presynaptic glutamate release due to P301L tau binding to synaptic vesicles to inhibit vesicle mobility and exocytosis.^6, 7^ More recent studies on the P301L tau mutation have shown that early in the disease process there is an increase in presynaptic glutamate release,^2^ and elevated synaptic activity called hyperexcitability,^3–5^ that is associated with memory deficits.^1^ The vast number of tau mediated changes in synaptic communication during disease progression do not occur in isolation and can also significantly affect other synaptic and neuronal functions. Thus, understanding this early-stage pathway of increased glutamate recycling is essential because it may lead to new early-stage treatments.

Using the rTg(TauP301L)4510 mouse model (hereafter called tau_P301L_ pos), we have previously shown that P301L tau expression increases glutamate release in the hippocampus, which correlated with memory deficits prior to observable neuron loss.^2^ We then showed that increased presynaptic glutamate release is directly caused by a tau-mediated increase in the number of vesicular glutamate transporter 1 (VGlut1) per glutamatergic vesicle.^8^ Recent studies have shown that tau mediates an increase in VGlut1 via altering nuclear epigenetic transcription factors.^9^ Increased VGlut1 per vesicle then increases the concentration of glutamate per vesicle,^8^ resulting in an increase in glutamate release per vesicle during activity.^2, 8,10^ Understanding the consequences of this molecular pathway is important because we separately showed that reducing VGlut1 levels and glutamate release rescued cognitive deficits in tau_P301L_ pos mice,^10^ suggesting the increased VGlut1 and glutamate release may represent potential therapeutic targets for the treatment of early-stage tauopathies.

Presynaptic vesicle release, recycling, and glutamate clearance are considered the predominant energy consumption processes in cognition.^11^ Further, excessive glutamate exposure has been shown to detrimentally affect mitochondria function, ^12^ as mitochondria dynamically respond to changes in neuronal and synaptic function. The density and structure of cristae within mitochondria rapidly and dynamically change in response to local ATP/ADP demand.^13, 14^ This change results in a change in the membrane potential,^15–17^ which is important because the potential difference across the mitochondria membrane drives ATP-synthesis.^13^ Equally important, mitochondria provide calcium buffering for local changes in synaptic activity.^16, 18^ Because VGlut1 is an ATP-dependent glutamate transporter and tau-mediates an increase in VGlut1 per vesicle, we hypothesize that local ATP demand will likely increase, resulting in increased demand from mitochondria.

It has been well established that reduced presynaptic function leading to neurodegeneration correlates with mitochondria dysfunction. Indeed, mitochondria dysfunction is a hallmark of tauopathy and neurodegeneration.^17, 19–22^ Mitochondria exhibit structural and morphological changes during AD.^23, 24^ As a consequence, mitochondria exhibit increased calcium adsorption,^18^ mitochondria-mediated neuronal apoptosis,^25^ and concomitant reduced presynaptic function. Additionally, presynaptic mitochondrial deficits are correlated with a presynaptic reduction in synaptic vesicles.^26^ However, with our recent observations of increased glutamate release prior to reduced presynaptic transmission, a central unanswered question is whether mitochondria morphology, structure, and function are altered in parallel with glutamate release during the early stages of disease progression, as hypothesized above.

It is also important to consider how mitochondria functional changes are either caused by, or support, P301L changes in synaptic function in order to better understand how early changes in mitochondria can lead to neurodegeneration. Mitochondria ATP synthesis utilizes a large combination of protein function and interactions,^27^ which break down with use and time,^28, 29^ and must be regularly replaced.^29^ Protein degradation was previously shown to lead to mitochondrial degeneration and ultimately neural degeneration in multiple different pathways.^30^ Separately, hyperexcitability is a pathway to increase trans-synaptic spread of tau,^3, 31^ and changes in mitochondrial function to support hyperexcitability thus represents a potential pathway to exacerbate disease progression. These possibilities warrant a careful study of mitochondria during early stage of tauopathy as a potential pathway to reduce or arrest disease progression.

In this present study, we sought to explore if mitochondria morphology and function are altered along with increased VGlut1 and glutamate release prior to observable presynaptic neurodegeneration. We use our previously established cell culturing approach of hippocampal neurons cultured at PND5 that showed increased VGlut1 and glutamate release.^8^ Here, we show that mitochondria morphology and structure change in P301L hippocampal neurons compared to hippocampal neurons from normal, non-transgenic litter mates (hereafter called tau_P301L_ neg). We show that altered structure correlates with decreased basal P301L membrane potential. We observe that post-activity mitochondria membrane potential remains higher in tau_P301L_ pos mitochondria compared to tau_P301L_ neg, and we show that reducing vesicle glutamate loading by inhibiting VGlut1 activity using Trypan Blue (TB) rescues differences in tau_P301L_ pos and tau_P301L_ neg mitochondria membrane potential after activity. Taken together, these results show for the first time that mitochondria dysfunction in a tauopathy model is mediated at least in part by increased glutamate release to support hyperexcitability.

## Materials and Methods

### Transgenic mouse colony and cell cultures

For this study, we used rTg4510 mice, a well-characterized and regulatable tauopathy mouse model of Alzheimer’s disease, which express human mutant P301L tau. The mice were generated by crossing a responder line with an activator line. The responder line contains cDNA of the human tau P301L mutation downstream of a tetracycline operon-responsive element (TRE) and the activator line contains a tetracycline-controlled transactivator (tTa) downstream of a Ca^2+^ calmodulin kinase II promoter. The bigenic progeny were genotyped for confirmation of the presence of both the activator and the promoter and wild type littermates were used as controls. All experiments conducted were compliant with the guidelines set forth by the International Animal Care and Use Committee and were approved by Auburn University Animal Care and Use Committee.

### La-SEM sample preparation and Analysis

Cells are prepared for La-SEM following our previously established protocol.^32^ Cell cultures were washed with cacodylate buffer (0.15 M cacodylate buffer and 2 mM CaCl2). Then, cells were aldehyde fixed for 15 minutes at 37°C in modified Karnovsky’s fixative (2.5% glutaraldehyde, 2% paraformaldehyde, and cacodylate buffer) and stored in fixative at 4 °C overnight. The next day, samples were washed with cacodylate buffer. A light-protected heavy metal incubation was then performed in a solution of 1.5% potassium ferrocyanide, 1% osmium tetroxide, and cacodylate buffer for 1 hour, followed by washing. Thiocarbohydrazide-osmium liganding (OTO) was performed via light-protected thiocarbohydrazide (aq) incubation at 60**°**C for 20 minutes, followed by a serial wash and a light-protected incubation in 2% osmium tetroxide (aq) for 30 minutes. Samples were washed and incubated in 1% uranyl acetate (aq) overnight at 4°C with light protection. The following day, samples were washed, and contrast was enhanced with *en bloc* staining in 20 mM lead aspartate at 60 **°**C for 30 minutes. Samples were again washed and underwent a stepwise dehydration series in ice-cold ethanol for 10 minutes each (Ethanol/ddH20: 20%, 50%, 70%, 90%, 100%, and 100%). After dehydration, samples underwent serial resin infiltration at 2-hour intervals (Durcupan/ethanol: 25%, 50%, and 75%). Samples were then placed in 100% Durcupan resin overnight and then into fresh 100% Durcupan resin with the accelerator. Cell coverslips are then placed on an aluminum weigh boat and cured in a 60 °C oven for 48 hours.

Post resin curing, samples were sent to the Washington University Center for Cellular Imaging (WUCCI) where the coverslips were exposed with a razor blade and etched off with concentrated hydrofluoric acid. Small pieces of the resin containing the cells were then cut out and mounted onto blank resin stubs before 50 nm thick five serial sections were cut in the cell culture growing plane and placed onto a silicon wafer chips. These chips were then adhered to SEM pins with carbon adhesive tabs and large areas (∼ 300 x 300 µm) were then imaged at high resolution in a FE-SEM (Zeiss Merlin, Oberkochen, Germany) using the ATLAS (Fibics, Ottowa, Canada) scan engine to tile large regions of interest. High-resolution tiles were captured at 16,384 x 16,384 pixels at 5 nm/pixel with a 5 µs dwell time and line average of 2. The SEM was operated at 8 KeV and 900 pA using the solid-state backscatter detector. Tiles were aligned using ATLAS 5 La-SEM images were analyzed using either imageJ or AMIRA-3D (thermofisher). Images were first smoothed using a 1-pixel gaussian blur rolling window. Mitochondria were chosen to satisfy the following restrictions: (1) The majority of the mitochondria outer membrane was intact; (2) The mitochondria was between 0.5 – 2 micron in length; (3) The mitochondria had at least six cristae to measure cristae distance; (4) The mitochondria was near observable microtubule axonal network. All of these restrictions limited the number of mitochondria per sample to ∼50 for both conditions.

Mitochondria cristae distance analysis was performed by obtaining line-intensity profiles for three cristae repetitions starting from a cristae peak location (Yellow lines, Fig. 2A) and normalizing to the starting intensity so that all data start at 1 (Fig. 2D). Each mitochondrion had at least two linescans and each linescan was counted equally. All line scans were averaged to obtain an aggregate correlation distribution of cristae intensities, and then fit to a sine function with decaying amplitude (Fig. 2E):

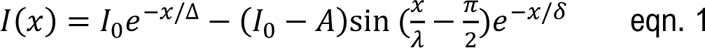

The distance to the first intensity peak was calculated from each linescan using a second-order derivative calculation. The average distance from the distribution of first cristae peak for each condition was then compared (Fig. 2F).

Calcium granules analysis was performed by counting the number of observable dark circles in La-SEM data that had at least 1.5x intensity above measured cristae intensity with a minimum size of 4 pixels in diameter. Calcium granule count was then divided by the total area of the mitochondria and aggregate results were compared between conditions (Fig. 3C).

Inner membrane fraction analysis was performed by thresholding individual mitochondria, pixel counts were either 1 for above threshold and 0 for below threshold, in La-SEM data with the restriction that the top 25% of intensity was above the threshold and the remainder of intensity was below (Fig. 3 B). Threshold images were then integrated with a boundary box inside the mitochondria (Dashed Red boundary in Fig. 3B), with the boundary set at 3-4 pixels inside the outer mitochondria membrane. The total number of pixels above threshold were then integrated. Aggregate distributions of total internal counts were compared for both conditions (Fig. 3 D).

Relative intensity data was performed by integrating the total amount of intensity (both inner and outer membrane) by drawing a boundary box around individual mitochondrion. Total integrated counts were then normalized by the area of the mitochondria to get counts per pixel. Integrated counts were also normalized by the average counts per cytosol to account for variability in background sample intensity. Aggregate normalized counts per pixel were then compared for both conditions (Fig. 3E).

### VGlut1-pHluorin transfection and imaging

VGlut1-pHluorin was generously provided by Drs. Robert Edward and Susan Voglmaier (UCSF).^33^ Lentiviral vectors were generated by the Viral Vectors Core at Washington University, and utilized in accordance with approved procedures at Auburn University. Cultures were transfected at DIV3 for 48 hrs. The media was then replaced with fresh neurobasal media, and half media replacements occurred every 3 days until day of imaging. During imaging, samples were exposed to imaging media containing 140 mM NaCl, 2.5 mM KCl, 2 mM CaCl_2_, 4 mM MgCl_2_, 10 mM HEPES, 2 mM Glucose, 50 mM DL-AP5, 10 mM CNQX, pH adjusted to pH 7.4, following our previously established protocol. Cells were imaged on a Nikon Ti-2 microscope (Nikon) using an EPI fluorescent GFP cube (Nikon) with a 100x Oil immersion objective (Nikon) and ORCA Flash v4 CMOS camera (Hamamatsu), with a final 0.065×0.065 μm pixel size, and temperature was maintained at 37C. Cells were then electrically field stimulated using two electrodes to depolarize the membrane and open calcium channels. Cells were stimulated using one of three protocols as described in each section.

### Spontaneous VGlut1-pHluorin Data analysis

Spontaneous VGlut1-pHluorin experiments were performed by first imaging cells in the absence of stimulation for 80 seconds, followed by 1Hz stimulation for 20 seconds, then 40Hz stimulation for 10 seconds, finally exposed to NH_4_Cl. Individual frames from VGlut1-pHluorin movies were background subtracted using a 10-frame rolling-ball-radius package in imageJ. Presynapses were then identified during 40Hz stimulation using in-lab matlab scripts. Identified presynapse location intensities were then run through a Kennedy-Chung filter algorithm previously shown to identify spontaneous release events.^34^ We then counted intensity spikes following our previously established approach of at least 1 standard deviation above noise level.^8^ The number of identified release events were then aggregated and divided by the observation time to estimate release frequency. The intensity per vesicle was determined as the three frame average of the peak minus the three-frame average of baseline before release.

### Low and High Activity VGlut1-pHluorin Data Analysis

Low and high activity VGlut1-pHluorin experiments were performed by first imaging cells in the absence of stimulation for 10 seconds, followed by a bout of stimulation (either 10Hz or 40 Hz for 10 seconds), then a delay for 10 seconds, followed by a second bout of stimulation 40Hz stimulation (either 10Hz or 40 Hz for 10 seconds), finally exposed to NH_4_Cl. Individual frames from VGlut1-pHluorin movies were background subtracted using a 30-frame rolling-ball-radius package in imageJ.

### MitoView 650 staining and imaging

At time of imaging, cell cultures were incubated in imaging media (See above) that contained 200 nM MitoView. Cultures were incubated for 15 minutes prior to imaging. Red fluorescence was measured using a DS-Red/TRITC/CY3 cube (Nikon). Each fluorescent image was taken with the same 100 msec exposure time. Images were analyzed using imageJ software tool sets. First, images were background subtracted using a 50-pixel rolling window. Second, images were smoothed using a 2-pixel gaussian blur rolling window. Line-scans were then taken (Fig. 1 A) along the long-axis and short-axis and the length/width were determined based on the full-width at half-maximum intensity of the images (Fig. 1 A).

**Fig 1:**
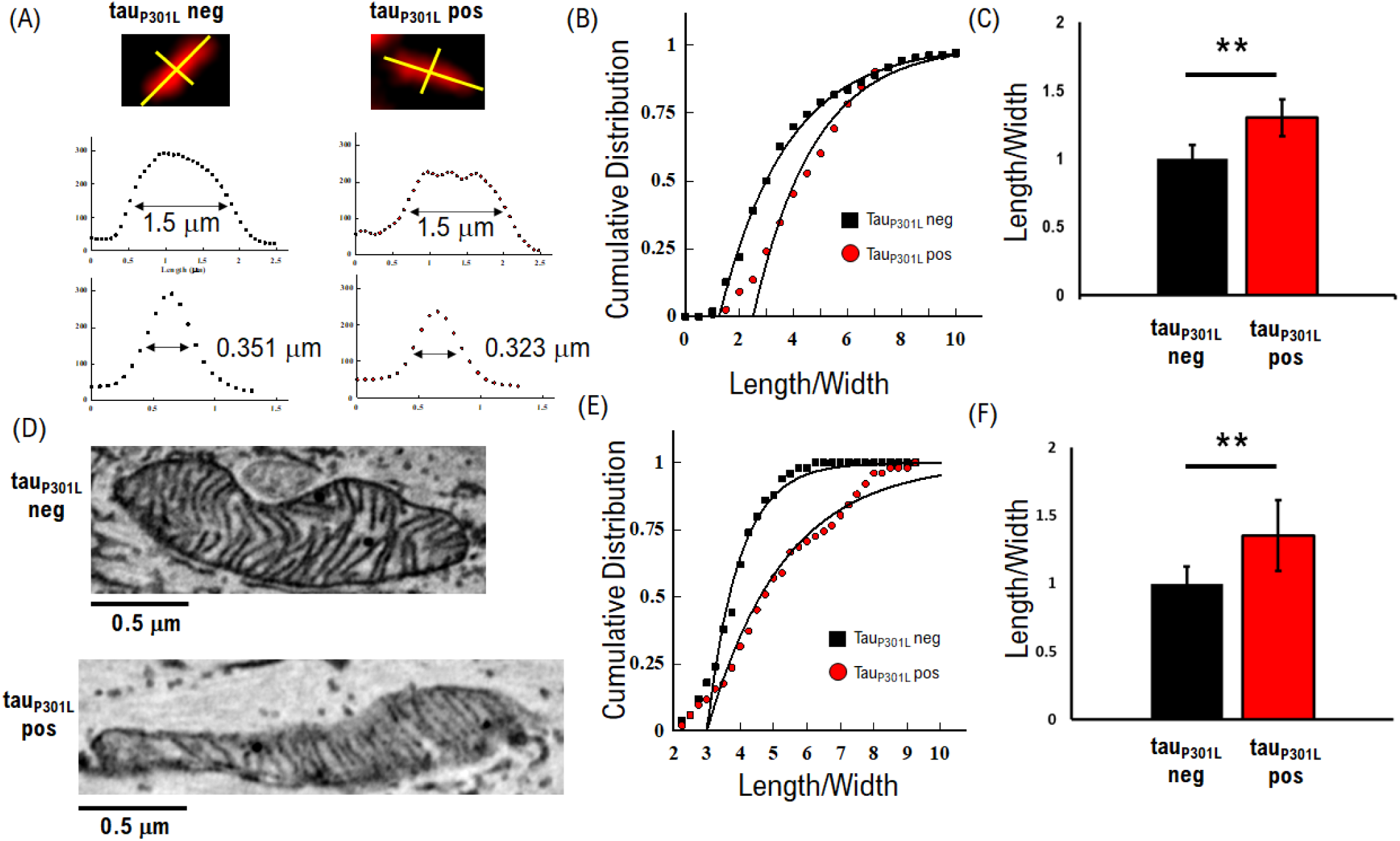
Baseline mitochondria structural differences in tau_P301L_ pos cultures versus tau_P301L_ neg cultures. (A) Example fluorescence images of individual mitochondria in tau_P301L_ neg (Top) tau_P301L_ pos cultures (Bottom). Measurements on length and width are obtained using line-scan intensity analysis. (B) The ratio of length/width shows tau_P301L_ pos mitochondria exhibit longer mitochondria than tau_P301L_ neg cultures. (C) Relative Length/Width ratio for FM results shows a 1.35 increase for tau_P301L_ pos (Red) mitochondria compared to tau_P301L_ neg cultures (Black). (D) Example La-SEM images of individual mitochondria in tau_P301L_ neg (Top) tau_P301L_ pos cultures (Bottom). Length and width of mitochondria are measured using line scan analysis (See Methods). (E) The ratio of length/width shows tau_P301L_ pos mitochondria exhibit longer mitochondria than tau_P301L_ neg cultures. The difference in length to width and overall scale are consistent with Fluorescence Microscopy results (B). (F) Relative Length/Width ratio for EM results shows the same 1.35 increase for tau_P301L_ pos (Red) mitochondria compared to tau_P301L_ neg cultures (Black), as observed for FM results (C). MitoView data: tauP301L neg: 110 mitochondria from 2 samples and 1 litter; tauP301L pos: 220 mitochondria from 4 samples and 2 litters; P-value from two-tailed t-Test. La-SEM data: tauP301L neg: 56 mitochondria from 1 sample and 1 litter; tauP301L pos: 100 mitochondria from 1 sample and 1 litter; P-value from two-tailed KS-Test. All error bars are SEM.

### JC-1 staining and imaging

At time of imaging, cell cultures were incubated in imaging media (See above) that contained 20 μg/mL JC-1, which is sufficient to form J-aggregates above the critical J-aggregate concentration.^35^ Media pH of 7.4 was used consistent with VGlut1-pHluorin imaging media and consistent with physiological conditions. Cultures were incubated with JC-1 media for 15 minutes prior to stimulation. Green fluorescent intensity was measured using a GFP cube (Nikon), and Red fluorescence was measured using a DS-Red/TRITC/CY3 cube (Nikon). The incoming excitation intensity was kept the same for both channels. Each fluorescent image was taken with the same 100 msec exposure time for both channels.

### JC-1 Fluorescent Image Data Analysis

JC-1 images were analyzed using imageJ software tool sets. First, images were background subtracted using a 50-pixel rolling window. Second, a mask was created using the red-channel image: (i) the background subtracted image was smoothing using a 2-pixel gaussian blur; (ii) the blurred imaged was thresholded so that the top 15-30% of intensity points were accepted; (iii) the imageJ analyze particle package was used to identify individual mitochondria consistent with the size observed in La-SEM data (0.2 – 4 μm^2^); (iv) an in-lab matlab script was written to integrate the intensity in the background-subtracted Red and Green images over all identified individual mitochondria; (v) a distribution of the ratio of Red/Green integrated intensities were binned and fit to normal and cumulative distributions. Mean values are reported from cumulative distributions and standard error of means are reported from normal distribution fits.

### RHOD-2 Fluorescent Image Data Analysis

RHOD-2 images were analyzed using imageJ software tool sets. First, images were background subtracted using a 50-pixel rolling window. Second, a mask was created using the red-channel image: (i) the background subtracted image was smoothing using a 2-pixel gaussian blur; (ii) the blurred imaged was thresholded so that the top 15-30% of intensity points were accepted; (iii) the imageJ analyze particle package was used to identify individual mitochondria consistent with the size observed in La-SEM data (0.2 – 4 μm^2^); (iv) an in-lab matlab script was written to integrate the intensity in the background-subtracted Red images over all identified individual mitochondria; Mean values are reported from cumulative distributions and standard error of means are reported from normal distribution fits.

### Computational Model

Computational simulations were based on our previously established binomial model,^8^ and based on previous models of presynaptic release ^36–38^. Here we performed multinomial vesicle release model of multiple release sites described as:

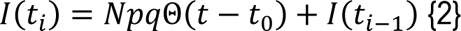

where N is the number of active zone sites, p is the probability of release per site (assumed to be constant for all sites), q is the intensity per vesicle. All model parameters used to reproduce each experimental conditions were constrained (Table S1). We also used a coarse-grained exponentially decaying vesicle release probability (*p*(*t*) = *p*_0_ + Δ*p* ∗ *e*^−*t*/*τp*^) to represent combined presynaptic factors that have been extensively explored but are beyond the scope of this present study. Simulations were performed using python 3.10 release, and the algorithm follows our previous approach.^8^

### Statistical Analysis

All data comparisons were supported by established statistical analysis appropriate for the experimental comparison. All VGlut1-pHluorin data were compared using Mann-Whitney U test. JC-1 comparisons of absolute differences were performed using two-tailed t-Tests. JC-1 comparisons on relative differences were performed using two-tailed KS-test. RHOD-2 comparisons of absolute differences were performed using two-tailed t-Test. All iGluSnFr comparisons were performed using Mann-Whitney U test. All L/W ratio data for MitoView 650 and La-SEM data were performed with two-tailed t-Test. Single vesicle VGlut1-pHluorin comparisons for release frequency and intensity were performed with two-tailed t-Tests.

## Results

### Tau_P301L_ pos Mitochondria Exhibit Morphological Changes

We first established basal mitochondria morphological and structural changes in tau_P310L_ pos compared to tau_P310L_ neg cultures. ATP production and calcium adsorption are proportional to overall mitochondrial size and morphology, where larger mitochondria with more cristae produce more ATP and buffer more calcium. To determine if P301L tau affects mitochondria, we measured individual mitochondria size using the fluorescent membrane marker MitoView 650, and our previously established Large Area Scanning Electron Microscopy (La-SEM) approaches.^32^ Here we used the length/width ratio to distinguish changes in morphology.

We observed that tau_P310L_ pos cultures exhibit changes in mitochondria morphology compared to tau_P310L_ neg cultures. We used a length/width analysis of MitoView 650 to quantify changes in morphology (Fig. 1 A) and observed that tau_P310L_ pos cultures show more narrow mitochondria than tau_P310L_ neg cultures (Fig. 1 A, B). To limit bias from differences in length, we limited the length distribution of measured mitochondria to less than 2 μm for both tau_P310L_ neg (mean length 1.87 +/- 0.09 μm) and tau_P310L_ pos (mean length 1.74 +/- 0.05 μm). Tau_P310L_ pos mitochondria had a larger **L/W** (4.4 +/- 0.133) compared to tau_P310L_ neg mitochondria (3.4 +/- 0.1). This increase is ∼30% increased relative ellipticity for tau_P310L_ pos mitochondria tau_P310L_ neg mitochondria (Fig. 1 C).

To support fluorescence microscopy results, we quantified mitochondria size in tau_P310L_ pos and tau_P310L_ neg cultures using La-SEM.^32^ We observed mitochondria in tau_P310L_ pos cultures exhibit increased size compared to tau_P310L_ neg cultures (Fig. 1 D). We used the same **L/W** ellipticity measure used for FM data, along with the same restriction in size distribution and found that tau_P310L_ pos mitochondria have a larger ratio (5.4 +/- 0.45, Fig. 1 E) than tau_P310L_ neg mitochondria (4.0 +/- 0.22, Fig. 1 E). Consistent with FM results, the increased ratio is due to a change in width rather than length. Further, tau_P310L_ pos mitochondria have the same increase in relative **L/W** compared to tau_P310L_ neg mitochondria (35%, Fig. 1 F, P = 0.0094, two-tailed KS-test) for La-SEM as observed for FM MitoView 650 (Fig. 1 C), providing further confirmation of a morphological change in mitochondria from tau_P310L_ pos cultures.

To determine the consequences of mitochondria morphological changes, we measured internal mitochondrial structural changes by measuring cristae density in individual mitochondria measured with La-SEM. Cristae are the functional locations of ATP production, and any changes in cristae structure will alter the dynamics of ATP turnover, particularly during presynaptic activity.^39, 40^ We measured cristae density in two ways: (i) measuring cristae diameter by quantifying the width from line-scan analysis of individual cristae observed in La-SEM data (Yellow lines, Fig. 2 A). (ii) by measuring nearest-neighbor distances between cristae; both measurements used line scan analysis of La-SEM images (Fig. 2 A, See Methods). We then compared aggregate cristae measurements to determine differences between tau_P310L_ pos and tau_P310L_ neg mitochondria.

**Fig 2:**
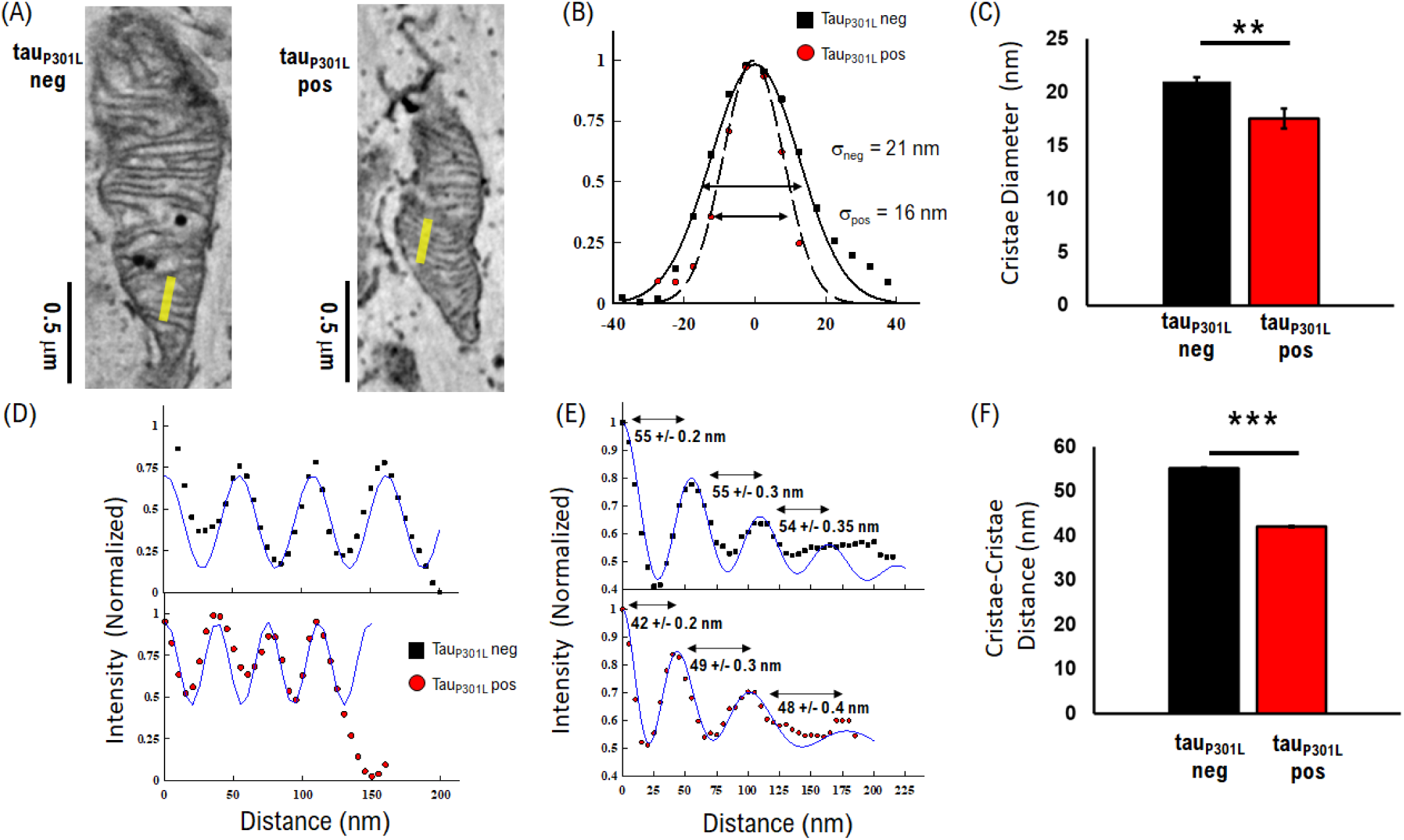
Baseline mitochondria Structural differences in tau_P301L_ pos cultures versus tau_P301L_ neg cultures. (A) Example La-SEM images of individual mitochondria in tau_P301L_ neg (Left) tau_P301L_ pos cultures (Right). (B) Example cristae width measured using line-scan analysis of La-SEM data. (C) Aggregate cristae width comparing tau_P301L_ neg (Black) and tau_P301L_ pos cultures (Red). (D) Distance between cristae is measured using line scan intensity analysis of EM data (E) Nearest-Neighbor intensities show a decrease in tau_P301L_ pos cultures. (F) Aggregate nearest-neighbor averages show a significant decrease in cristae distance for tau_P301L_ pos (Red) mitochondria compared to tau_P301L_ neg cultures (Black). Cristae Width: tauP301L neg: 42 cristae from 1 sample and 1 litter; tauP301L pos: 41 cristae from 1 sample and 1 litter; P-value from two-tailed t-Test Cristae Distance: tauP301L neg: 129 scans from 1 sample and 1 litter; tauP301L pos: 109 scans from 1 sample and 1 litter; P-value from two-tailed t-Test. All error bars are SEM.

Cristae diameter decreased significantly for tau_P310L_ pos mitochondria compared to tau_P310L_ neg. We observed that tau_P310L_ pos mitochondria were typically more narrow (Red circles, Fig. 2 B) than tau_P310L_ neg mitochondria (Black squares, Fig. 2 B). Averaging individual cristae across multiple mitochondria shows that tau_P310L_ pos cristae diameter (17.5 +/- 0.9 nm) was significantly smaller (P = 0.002, 2-tailed t-test) compared to tau_P310L_ neg cristae (21 +/- 0.37 nm). Further, tau_P310L_ pos cristae diameter exhibited more variability than tau_P310L_ neg cristae, as measured by the increased error for the same number of analyzed cristae (tau_P310L_ pos for 0.9 nm SEM, compared to 0.37 SEM for tau_P310L_ neg).

We observed that tau_P310L_ pos cultures exhibit an increased density of cristae compared to tau_P310L_ neg cultures (Fig. 2 A). Mitochondria from both tau_P310L_ pos and tau_P310L_ neg cultures exhibited a regular periodic distribution of cristae throughout the internal compartment (Fig. 2 A), consistent with previously observed hippocampal neuronal mitochondria.^13, 41, 42^ However, tau_P310L_ pos mitochondria exhibited cristae with a shorter periodicity (Fig. 2 D) compared to tau_P310L_ neg cultures (Fig. 2 B). Aggregate tau_P310L_ neg mitochondria showed nearest-neighbor cristae periodicity of 55 +/- nm (Fig. 2 C), compared to 40 +/- nm for tau_P310L_ pos mitochondria (Fig. 2 C). Further, tau_P310L_ pos mitochondria exhibited more variability in cristae distance with average nearest neighbor distances initially 40 nm apart (Red Circles, Fig. 2 C), but the distance between neighboring cristae increases to 50 nm as the distance increases. In contrast, the distance between neighboring cristae remains a constant 50 nm for tau_P310L_ neg mitochondria (Black Squares, Fig. 2 C).

We next established if altered cristae structure resulted in calcium adsorption or more total membrane within the mitochondria under resting conditions. We used an integrated intensity analysis approach to measure changes in mitochondria membrane material (Fig. 3 A), which includes proteins used for ATP production and adsorbed material.^43^ We then used a masking approach to determine the amount of membrane-occupied surface per unit area (Fig. 3 B), which can affect the amount of membrane used for ATP production.^13^ Finally, we measured adsorbed calcium by quantifying the number of calcium granules per unit area (Fig. 3 A).

**Fig 3:**
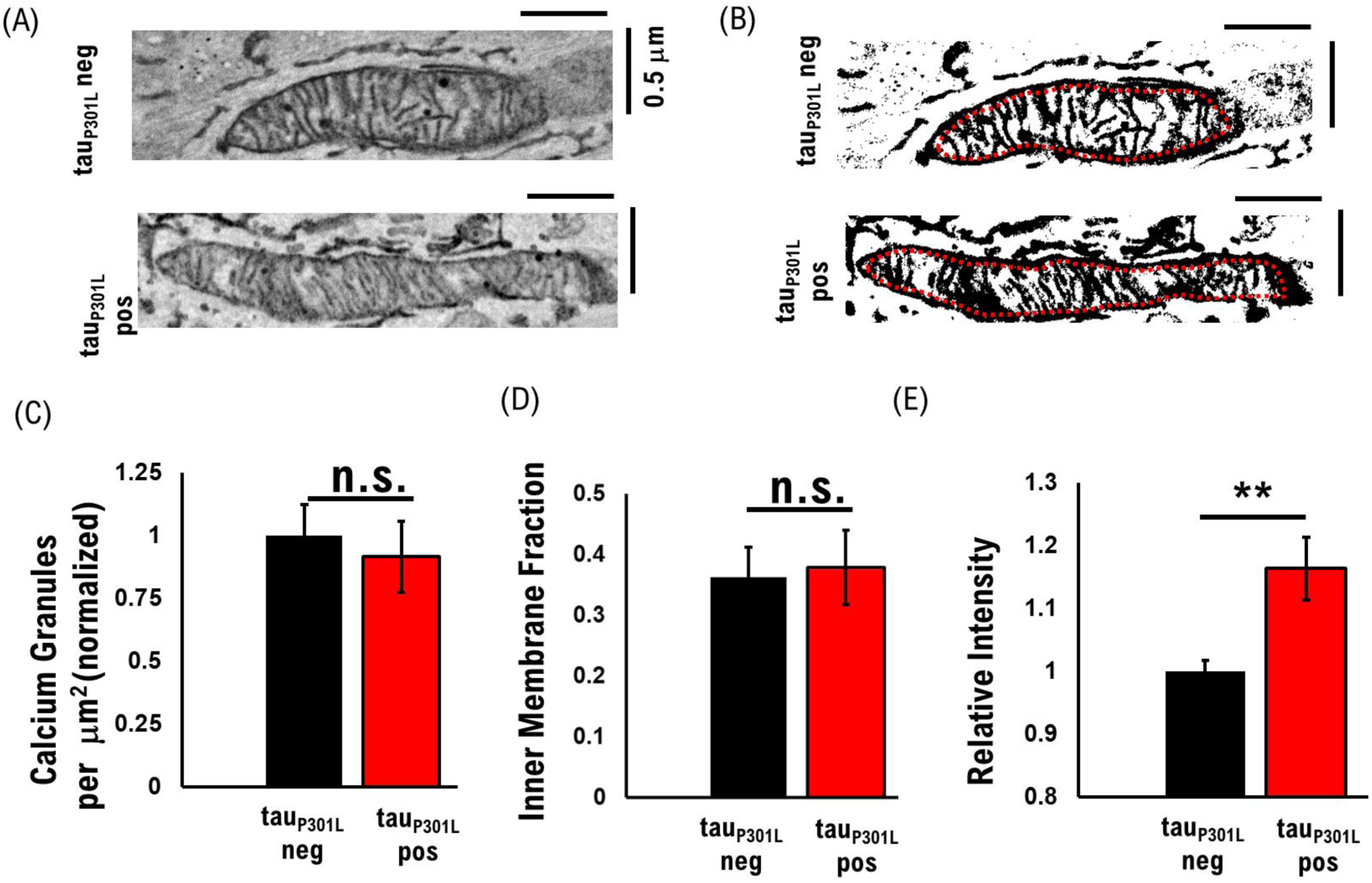
Baseline mitochondria calcium adsorption and membrane density are unchanged while the amount of membrane material increases for tauP301L pos neurons. (A) Example EM imaged mitochondria with adsorbed calcium granules and membrane intensity (B) Example integrated area analysis of masked mitochondria. (C) The density of Calcium granules is unchanged for tau_P310L_ neg (black) and tau_P310L_ pos (red) mitochondria. (D) The total amount of inner membrane material is unchanged for tau_P310L_ neg (black) and tau_P310L_ pos (red) mitochondria. (E) The membrane intensity increases for tau_P310L_ pos (red) mitochondria compared to tau_P310L_ neg (black). tau_P301L_ neg N = 50 from 1 sample from 1 litter; tau_P301L_ pos N = 49 from 1 sample from 1 litter; P = 0.0233, two-tailed t-Test

We observed that the amount of adsorbed calcium in calcium granules per unit area is unchanged for tau_P310L_ pos (0.92 +/- 0.14) and tau_P310L_ neg (1.0 +/- 0.12) under resting conditions (Fig. 3 C). We also observed that the total fraction of the inner membrane area (Fig. 3 D) is the same for tau_P310L_ pos (0.38 +/- 0.06) and tau_P310L_ neg (0.36 +/- 0.05). However, we observed a significant increase (P < 0.01, two-tailed tTest) in the amount of intensity per unit area for tau_P310L_ pos (1.16 +/- 0.05) and tau_P310L_ neg (1.0 +/- 0.02).

Taken together, our La-SEM results show that the total mitochondria membrane is conserved for tau_P310L_ pos mitochondria compared to tau_P310L_ neg, however, the membrane surface area within the mitochondria is increased as a consequence of smaller cristae that are more densely packed. Further, the amount of metal-stained material within the membrane increases in tau_P310L_ pos mitochondria compared to tau_P310L_ neg. These results suggest that mitochondria morphology and structure are altered to support increased energy demands.

### Tau_P301L_ pos Mitochondria Maintain Elevated Membrane Potential After Activity

We next sought to establish the effects of tau_P301L_ on mitochondria membrane potential in basal conditions, because membrane potential drives production of ATP in order to support normal synaptic function.^16^ We used the established membrane potential reporter JC-1 to measure membrane potential. JC-1 absorbs into the inner mitochondria membrane as monomers that fluoresce in the green spectra at low mitochondria membrane potential and begin to form aggregates that fluoresce in the red spectra at higher membrane potentials.^35, 44^ The relative difference in red/green intensity changes linearly with the change in membrane potential above the critical J-aggregate concentration (CJC) between 30 mV – 200 mV.^35^ Consequently, JC-1 Red/Green intensity ratio changes correspond with mitochondria membrane potential.

We first confirmed that our approach identifies neuronal mitochondria by co-localizing identified mitochondria in the CY5 channel with processes observed using transmitted light (Fig. 4 A). This showed that the majority of all CY5 identified puncta co-localized with an observable neuronal process and that changes in mitochondria membrane potential reflected changes in neuronal mitochondria. To determine Red/Green fluorescence ratios, we then integrated the intensity of mitochondria in the Cy5 channel (yellow arrows, Fig. 4 A), and integrated the same identified location in the GFP channel (yellow arrows, Fig. 4 A), followed by quantifying aggregate results (See Methods).

**Figure 4:**
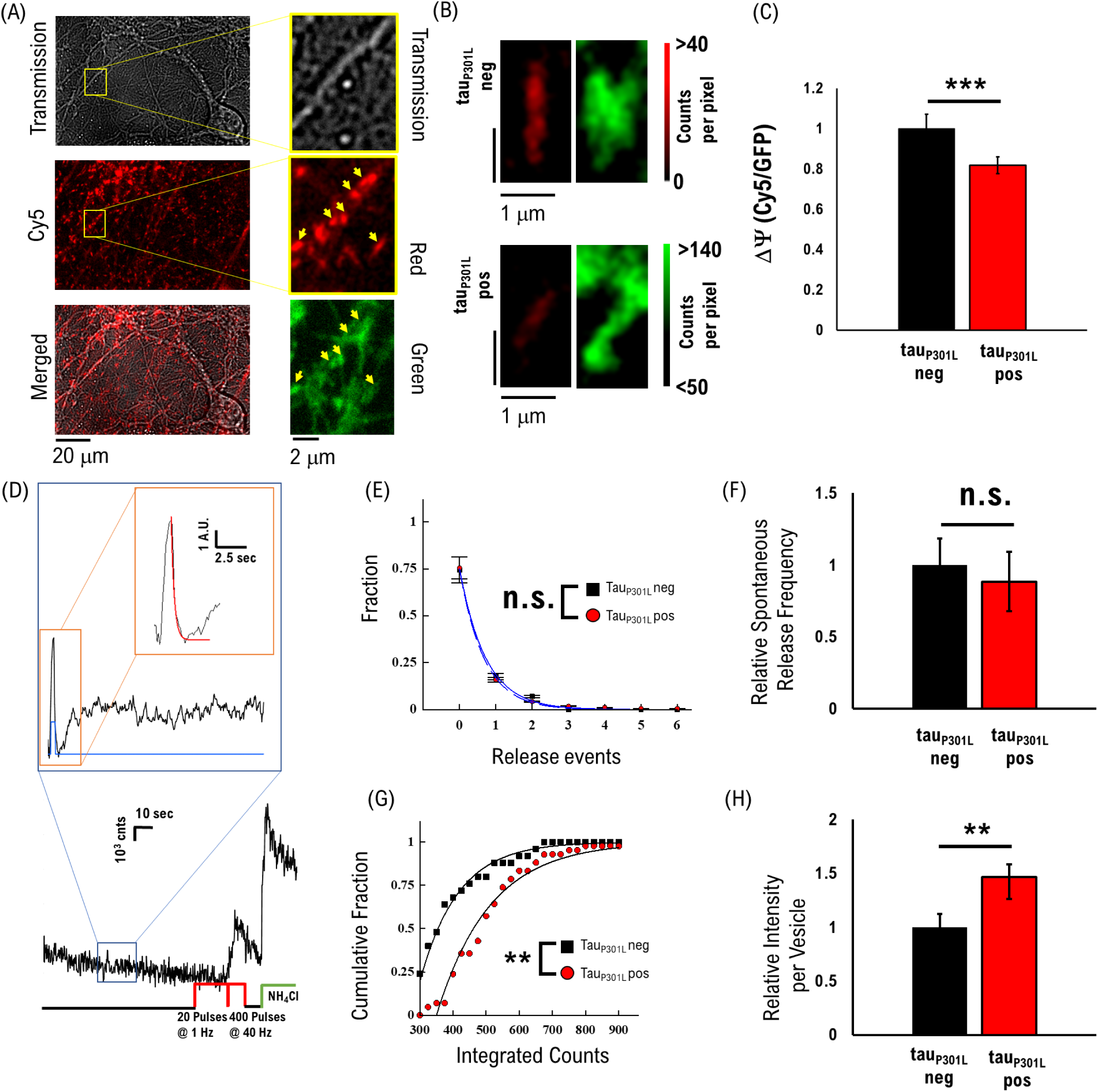
Baseline mitochondria membrane potential is lower for tauP301L pos neurons. (A) Example of cell cultures (Transmission Top Panel) with JC-1 stained mitochondria (Cy5, Middle Panel) and an overlay (Bottom Panel). Mitochondria are identified (yellow arrows) in the Cy5 channel (Cy5, Middle Right Panel) and co-localized with GFP intensity (GFP, Bottom Right Panel). (B) Example JC-1 stained mitochondria from tau_P310L_ neg (left) and tau_P310L_ pos (right) cultures. Tau_P310L_ pos mitochondria exhibit lower Cy5 relative to GFP intensity than tau_P310L_ neg neurons. (C) Aggregate intensity ratio for tau_P310L_ neg (black) and tau_P310L_ pos (red) normalized to tau_P310L_ neg. Tau_P310L_ pos mitochondria intensity ratio is 20% lower than tau_P310L_ neg. (D) Example experimental measurement including a single spontaneous release trace for tau_P310L_ pos presynapse (F) Fraction number of observed release events for tau_P310L_ neg (black) and tau_P310L_ pos (red) per minute. (G) Aggregate spontaneous release frequency for tau_P310L_ neg (black) and tau_P310L_ pos (red) normalized to tau_P310L_ neg. (H) Cumulative integrated intensity per vesicle for tau_P310L_ neg (black) and tau_P310L_ pos (red) normalized to tau_P310L_ neg. (I) Aggregate intensity per vesicle for tau_P310L_ neg (black) and tau_P310L_ pos (red) normalized to tau_P310L_ neg. Tau_P310L_ pos vesicle intensity is 40% higher than tau_P310L_ neg.

We first observed that the JC-1 stained mitochondria (Fig. 4 B) exhibit lower intensity in the CY5 channel for tau_P310L_ pos compared to tau_P310L_ neg cultures, indicating reduced JC-1 aggregates and thus lower membrane potential. We then quantified aggregate differences in intensity and observed that tau_P310L_ pos mitochondria exhibited a significant reduction (0.86 +/- 0.0375) intensity ratio (Fig. 4 C, P <0.001) compared to tau_P310L_ neg mitochondria (1.0 +/- .007). This ∼15% reduction in intensity ratio would correspond to an ∼ 20 mV reduction in membrane potential measured by JC-1.^35^ This difference suggests that baseline mitochondria potential correlates with differences in mitochondrial morphology and structure.

Since homeostatic mitochondria membrane potential reflects differences in homeostatic synaptic function, we determined if there were any differences in the presynaptic homeostatic function that would support mitochondrial membrane potential differences. Hippocampal presynapses spontaneously release vesicles to maintain synaptic connections at a rate of ∼0.5 vesicles per second.^45^ Changes in the spontaneous release rate could thus be a mediating factor in baseline mitochondria membrane potential. We measured single vesicle release events using the VGlut1-pHluorin approach in the absence of stimulation (Fig. 4 D) followed by 40 Hz stimulation to identify presynaptic locations. We observed the majority of presynapses (∼74%) release no vesicles within the 80-sec time-period of observation for both tau_P310L_ pos and tau_P310L_ neg cultures (Fig. 4 E), consistent with a low probability of release events. Comparing aggregate spontaneous release measured from multiple cultures shows the same rate for both tau_P310L_ pos (0.477 +/- 0.1) and tau_P310L_ neg (0.54 +/- 0.1), which is not significantly different (Fig. 4 F). These results suggest that the frequency of spontaneous vesicle release does not contribute to differences in observed mitochondrial membrane potential.

While the rate of spontaneous release is unchanged, the number of vesicles per release event (called multi-vesicular release MVR)^46^ and/or amount of VGlut1 per vesicle could contribute to differences. We have previously shown that tau_P310L_ pos neurons exhibit increased VGlut1 per vesicle during stimulated release by quantifying integrated intensity per vesicle.^8^ We used the same approach to distinguish changes in aggregate VGlut1 intensity per vesicle during spontaneous release (Fig. 4G). We observed that aggregate vesicle intensity (Fig. 4 G,H) increases in tau_P310L_ pos vesicles (481 +/- 97 counts) compared to tau_P310L_ neg vesicles (329 +/- counts 40). This increased intensity per vesicle is the same (∼1.4x) intensity increase per vesicle as observed for stimulated released vesicles we previously measured.^8^ Further, the single-peak fit to both distributions (Solid lines Fig. 4G) suggests that only one vesicle is released for the majority of events for both tau_P310L_ pos and tau_P310L_ neg cultures.

These results show that the number and frequency of spontaneous release events do not contribute to changes in baseline mitochondria morphology, structure, and function, but increased VGlut1 per vesicle is a likely contributing factor to differences in mitochondria membrane potential. Further, altered mitochondrial morphology and function occur prior to any observable changes in presynaptic release mechanics, suggesting that these are early-stage changes.

### Activity-dependent Changes in Membrane Potential is Lower for Tau_P301L_ pos Mitochondria

To understand the functional consequences of altered mitochondria morphology and structure on active presynaptic function, we next measured the effects of presynaptic activity on mitochondria membrane potential as a function of time after activity. Presynaptic activity effects on mitochondria are essential because presynaptic vesicle release and recycling utilizes the majority of presynaptic ATP at every stage of the recycling process.^11, 47^ The use of ATP would thus require mitochondrial-based ATP production to replenish local ATP usage.^39, 40, 48^ To establish if P301L tau mitochondria morphology and membrane potential are altered under activity-dependent presynaptic function, we used the same JC-1 imaging reporter and quantified changes in membrane potential as a function of time after presynaptic activity (See Methods).

We used two different stimulation protocols to test the different physiological limits of hippocampal neuronal function. To test normal hippocampal activity, we used a lower stimulation protocol (100 pulses at 10Hz, Fig. 5 A), which releases a small fraction of the recycling pool and represents relatively normal ATP demand.^49^ This low stimulation is consistent with observed spontaneous firing rates,^50^ or an action-potential train during normal hippocampal function.^51^ To test hyperexcitable or seizure-like states observed in AD, which corresponds with an extended level of presynaptic release at a frequency greater than 20 Hz,^3, 50^ we used a high stimulation protocol (two bouts of 400 pulses at 40 Hz stimulation, Fig. 5 B). This extended bout of stimulation also depletes the majority of releasable vesicles and would thus represent a large ATP demand. Consequently, these two protocols test the role of presynaptic release mechanics during normal neuronal function and hyperexcitable states, as well as their consequences on mitochondrial function.

**Fig 5:**
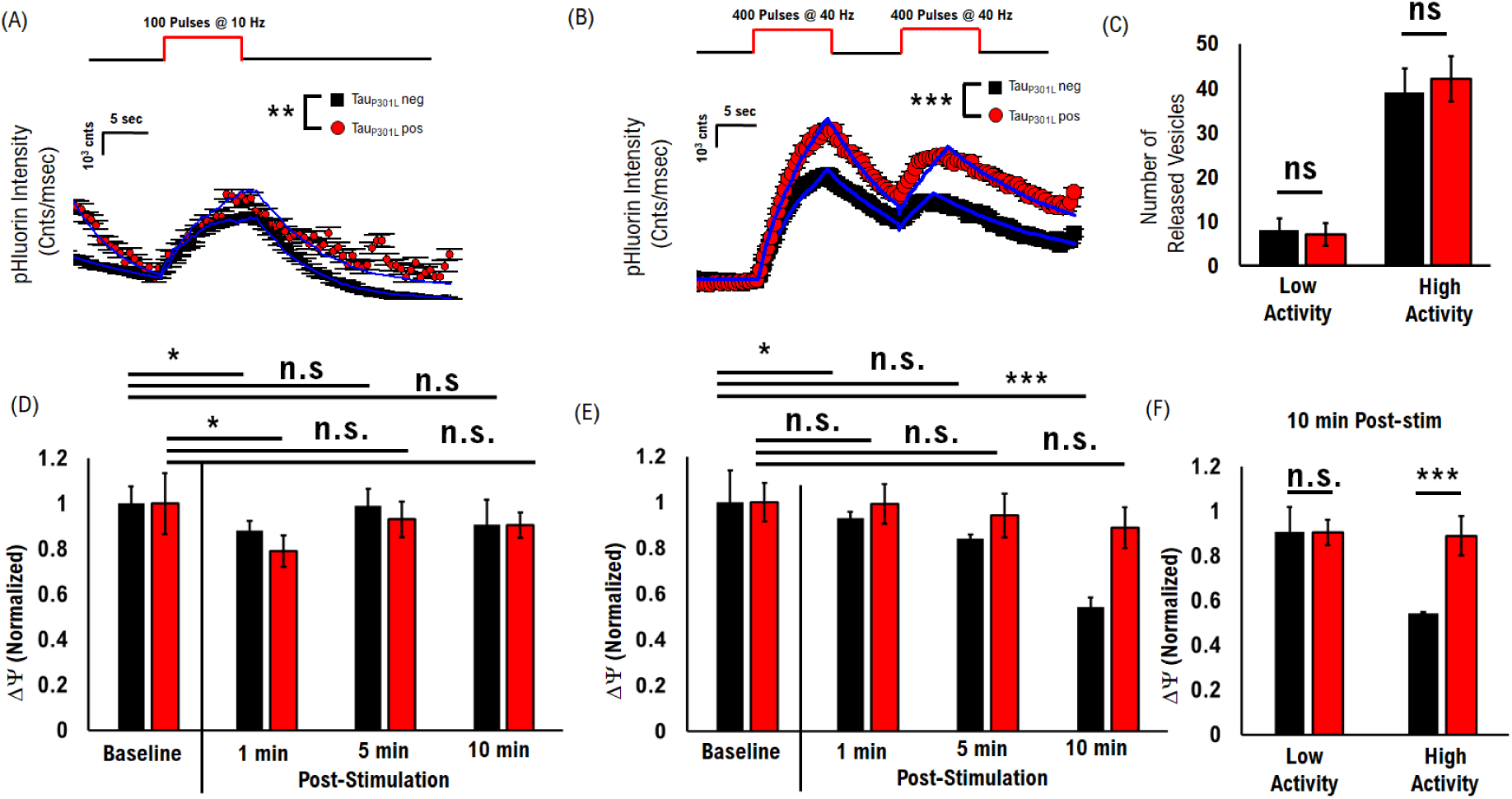
Activity-dependent changes in Mitochondria Membrane potential in tau_P301L_ pos neurons. (A) Presynaptic Response to Low Stimulation Protocol (100 Pulses at 10 Hz). Tau_P310L_ pos neurons show more VGlut1-pHluorin intensity than tau_P310L_ neg neurons, consistent with previously published results. Computational simulations (Blue lines) reproduce VGLUT1-pHluorin intensity based on a probabilistic model with intensity per vesicle as a free parameter (See Methods). (B) Presynaptic Response to High Stimulation Protocol (2 bouts of 400 stimulations at 40 Hz). Computational simulations (Blue lines) reproduce VGLUT1-pHluorin intensity based on a probabilistic model with intensity per vesicle as a free parameter (See Methods). (C) Number of released vesicles per low and high activity conditions are unchanged for Tau_P310L_ pos (Red) and Tau_P310L_ neg (Black) neurons. (D) JC-1 measured mitochondria membrane potential under Low Stimulation shows an initial decrease 1 min after stimulation, followed by a return to baseline for both Tau_P310L_ pos (Red) and Tau_P310L_ neg (Black) neurons. (E) JC-1 measured mitochondria membrane potential under High Stimulation shows a significant and sustained decrease for Tau_P310L_ neg (Black) neurons after stimulation, but a less substantial reduction for Tau_P310L_ pos (Red). (F) Relative JC-1 measured membrane potential 10 min after stimulation for low and high activity conditions, normalized to their respective baseline value. Low-stim: N>1000 mitochondria per sample from 2 plates and 1 litter for all conditions. High-stim: N>1000 mitochondria per sample from 5 plates and 2 litter for all conditions. *= P < 0.06; ** = P<0.01; *** = P < 0.001; pHluorin statistical comparisons are Mann-Whitney U test. JC-1 comparisons within rTg conditions are two-tailed KS-test of cumulative distributions. JC-1 comparisons across rTg conditions are pair-wise two-tailed t-Test.

Following our previously established VGlut1-pHluorin protocol,^8^ we observed a significant increase in VGlut1-pHluorin intensity for tau_P310L_ pos compared to tau_P310L_ neg cultures for both protocols, indicating an increase in the number of VGlut1 transporters per vesicle.^8^ We used our computational model approach^8^ to reproduce observed VGlut1-pHluorin intensity (blue lines in Fig. 5 A-B, See Methods) and determine the release probability and number of released vesicles. We found that both the release probability per stimulation and the number of released vesicles are unchanged for tau_P310L_ pos neurons compared to tau_P310L_ neg neurons (Fig. 5 C). These results show that the number of released vesicles is the same and thus does not contribute to any changes in mitochondria membrane potential under the same stimulation conditions.

We then measured JC-1 intensity ratio as a function of time after stimulation under both the low stimulation (**Fig. 5 D**) and high stimulation (**Fig. 5 E**). We observed that mitochondria membrane potential initially decreased after stimulation for the low stimulation protocol, followed by a return to baseline 10 min after stimulation for both tau_P310L_ pos and tau_P310L_ neg mitochondria (**Fig. 5 D**). However, the membrane potential significantly decreased as a function of time under the high stimulation protocol for tau_P310L_ neg mitochondria (**Black Fig. 5 E**); whereas tau_P310L_ pos mitochondria membrane potential remained constant or only slightly decreased (**Red Fig. 5 E**). At 10 min after stimulation, tau_P310L_ pos mitochondria were lower than baseline (0.78 +/- .11%), but not significantly different for the low stimulation or the high stimulation protocols (**Red Fig. 5 F**). The tau_P310L_ neg mitochondria exhibit a significant reduction under high stimulation (0.54 +/- 0.02%) at 10 min compared to baseline and compared to the low stimulation protocol (**Black Fig. 5 F**). Consequently, tau_P310L_ pos mitochondria membrane potential was 40% higher than tau_P310L_ neg mitochondria 10 min after stimulation (**Fig. 5 F**). It is important to note that mitochondria membrane potential returned to baseline 40-50 min after the high-activity protocol for both tau_P310L_ pos and tau_P310L_ neg mitochondria (**Fig. S2 F**).

To put the changes in JC-1 intensity ratio after high-activity in the context of mitochondria membrane potential, we compared the 10 min intensity ratio to the baseline tau_P310L_ neg mitochondria intensity. We observe that the tau_P310L_ pos mitochondria intensity ratio is ∼45% higher (DF/F = 0.89 +/- .09) than the tau_P310L_ neg mitochondria (DF/F = 0.54 +/- .04). This would correspond to a ∼12 mV greater membrane potential for tau_P310L_ pos mitochondria compared to tau_P310L_ neg mitochondria. These results suggest that tau_P310L_ pos mitochondria membrane potential changes are mitigated during high activity compared to tau_P310L_ neg mitochondria in order to support higher activity that occurs during hyperexcitable states.

We also note that the changes we observe here are independent of the specific P301L mouse model. We performed the same high-activity VGlut1-pHluorin experiments as well as the same JC-1 membrane potential measurements using cultures from the 3xTg mouse model that exhibits both beta-amyloid and tau pathology due to its three mutations associated with familial Alzheimer’s disease (APP Swedish, MAPT P301L, and PSEN1 M146V) We found an elevated VGlut1 level (**Fig. S1 B**) similar to the P301L tau mice, and the same time-dependent membrane potential after a bout of high activity (**Fig. S1 C-E**). These results support the hypothesis that increased VGlut1 mediates a change in mitochondria structure and function independent of the mouse model studied.

### Hypothesized Pathways for Observed Differences in Membrane Potential in Tau_P301L_ pos Mitochondria

To develop a better understanding of both the contributing factors to, and consequences of, observed differences in mitochondrial membrane potential, it is important to put the changes in JC-1 measured intensities in context of absolute membrane potential differences. To do this, we used previously observed JC-1 calibration curves of mitochondrial membrane potential,^35^ in combination with the known hippocampal neuronal mitochondria resting potential of ∼ −138 +/- 4 mV.^16, 35^ We then compared relative changes in membrane potential for both tau_P310L_ neg and tau_P310L_ pos mitochondrial (Fig. 6 A).

**Fig 6:**
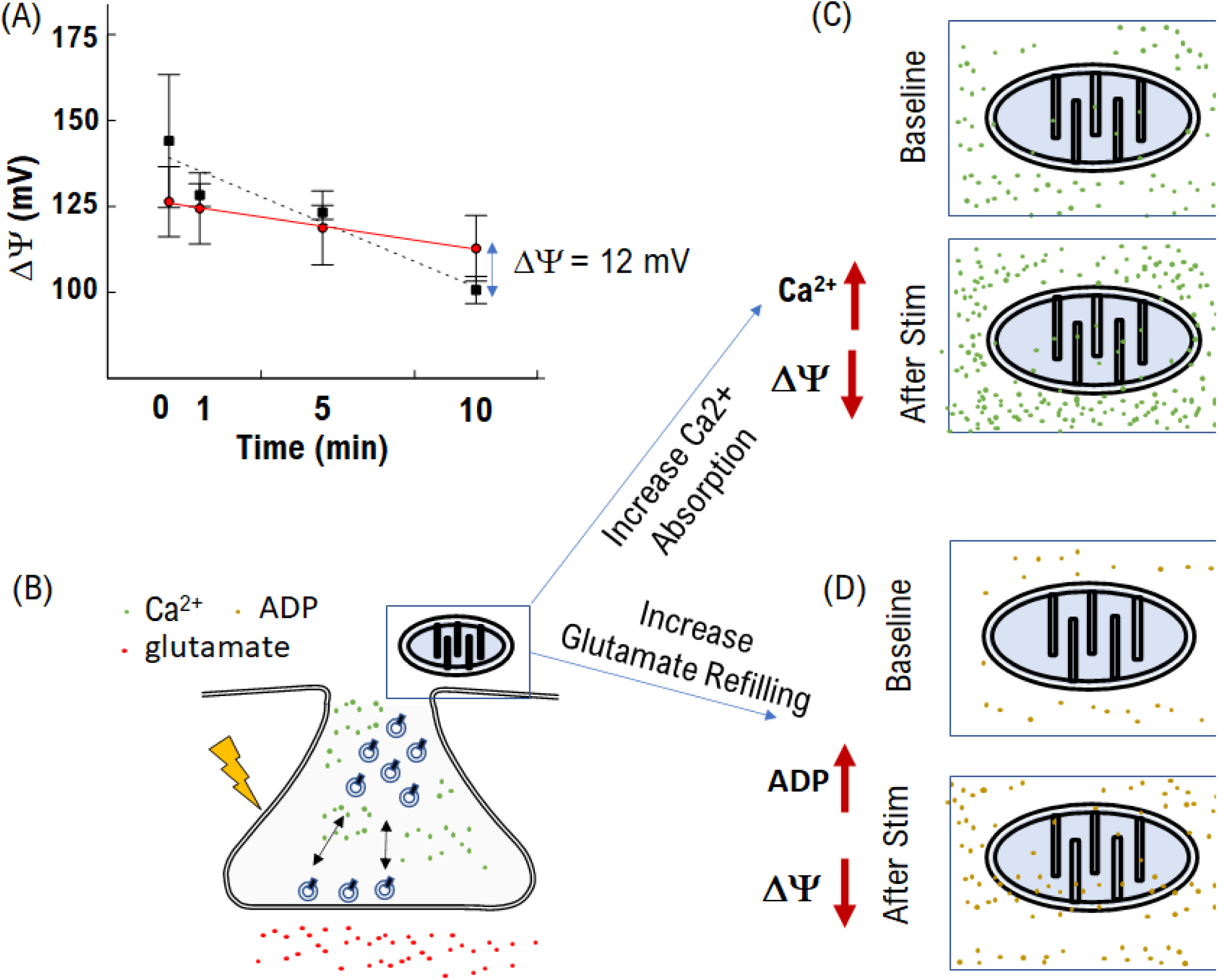
Hypothesized pathways to decreased membrane potential. (A) Mitochondria Membrane potential inferred from JC-1 intensity ratio (Fig. 5G) (B) Activity-dependent pathways to changes in mitochondria membrane potential (C) A pathway is by changes in the absorption of calcium after high activity (D) Another potential pathway to an activity-dependent membrane potential reduction is increased ATP production demand.

The baseline difference in JC-1 intensity for tau_P310L_ pos mitochondria would then correspond to a resting membrane potential of −122 +/- 10 mV (Fig. 6A). The membrane potential then decreases linearly with time after the bout of high activity (Fig. 6A), but the decrease is greater for tau_P310L_ neg (Dashed line, 3.7 mV/min, Fig. 6A) compared to tau_P310L_ pos (Solid line, 1.3 mV/min, Fig. 6A) mitochondria. The different time-dependent slopes in membrane potential result in tau_P310L_ pos mitochondria having a −12mV greater in membrane potential than tau_P310L_ neg mitochondria. This difference is crucial because it has been shown that every 10 mV difference in mitochondria membrane potential results in double the amount of ATP production.^52, 53^ Therefore, understanding the mechanisms that lead to the observed activity-dependent change in membrane potential is essential to understanding the physiological effects on neuronal function during AD progression.

Here we consider two main pathways to explain the difference in activity-dependent membrane potential observed after the bout of high-activity (**Fig 6 B-D**):

The first pathway involves the mitochondria functioning as a calcium buffer and absorbing calcium after activity (**Fig. 6C**). Presynaptic vesicle release involves a large influx of calcium for vesicle exocytosis and recycling.^54^ This influx must then be cleared to limit persistent vesicle release and prepare for the next evoked release event.^54^ Mitochondria absorption of calcium results in a reduction in membrane potential.^16^ Mitochondria act as calcium buffers by absorbing calcium both as granules and in the membrane.^55^ While we did not observe any difference in the concentration of calcium granules in tau_P310L_ pos mitochondria compared to tau_P310L_ neg mitochondria (**Fig. 3 C**), we did observe a significant difference in the membrane intensity using La-SEM that indicates an increase in the amount of heavy-metal stained membrane material. Suppose tauP310L pos mitochondria have more membrane-absorbed calcium at baseline, which would support the lower baseline membrane potential observed in tauP310L pos mitochondria compared to tauP310L neg mitochondria (**Fig. 4 C**). In that case, tau_P310L_ pos mitochondria may also have a reduced capacity for absorbing calcium after activity resulting in a lower activity-dependent change observed (**Fig 5 H**).

The second pathway involves ATP production (**Fig. 6D**). Presynaptic activity results in a large ATP use, significantly decreasing local ATP concentration.^56^ The drop in ATP concentration and increase in ADP results in a larger production of mitochondrial ATP, which in turn reduces membrane potential via the TCA cycle.^27, 57, 58^ If tau_P310L_ pos mitochondria have already restructured to support increased glutamate refilling during baseline, they may also provide increased ATP production after activity. Consequently, a change in activity-dependent membrane potential would be observed differently in tau_P310L_ pos mitochondria compared to tau_P310L_ neg mitochondria.

We will test the possible contribution of each pathway for the remainder of this study to determine if either (or both) pathways contribute to observed differences in activity-dependent membrane potential.

### Activity-Mediated Calcium Absorption is Unchanged in Tau_P301L_ pos Mitochondria

To understand if the pathway of calcium absorption (**Fig. 6D**) mediates the observed changing mitochondria membrane potential, we measured adsorbed calcium after the high stimulation protocol using the established RHOD-2 reporter. RHOD-2 fluorescence intensity is proportional to the amount of calcium adsorbed in mitochondria.^59^ We followed the same staining protocol used for JC-1 by first identifying RHOD-2 puncta along observed neuronal processes (**Fig. 7 B**). We then integrated intensity of observed RHOD-2 puncta and compared aggregate mean intensity differences (See Methods). Finally, we normalized all intensities to the baseline tau_P301L_ neg level. If tau_P301L_ pos mitochondria exhibit significant differences in absorbed calcium, then tau_P301L_ pos mitochondria will have elevated RHOD-2 intensity compared to tau_P301L_ neg mitochondria.

**Fig 7:**
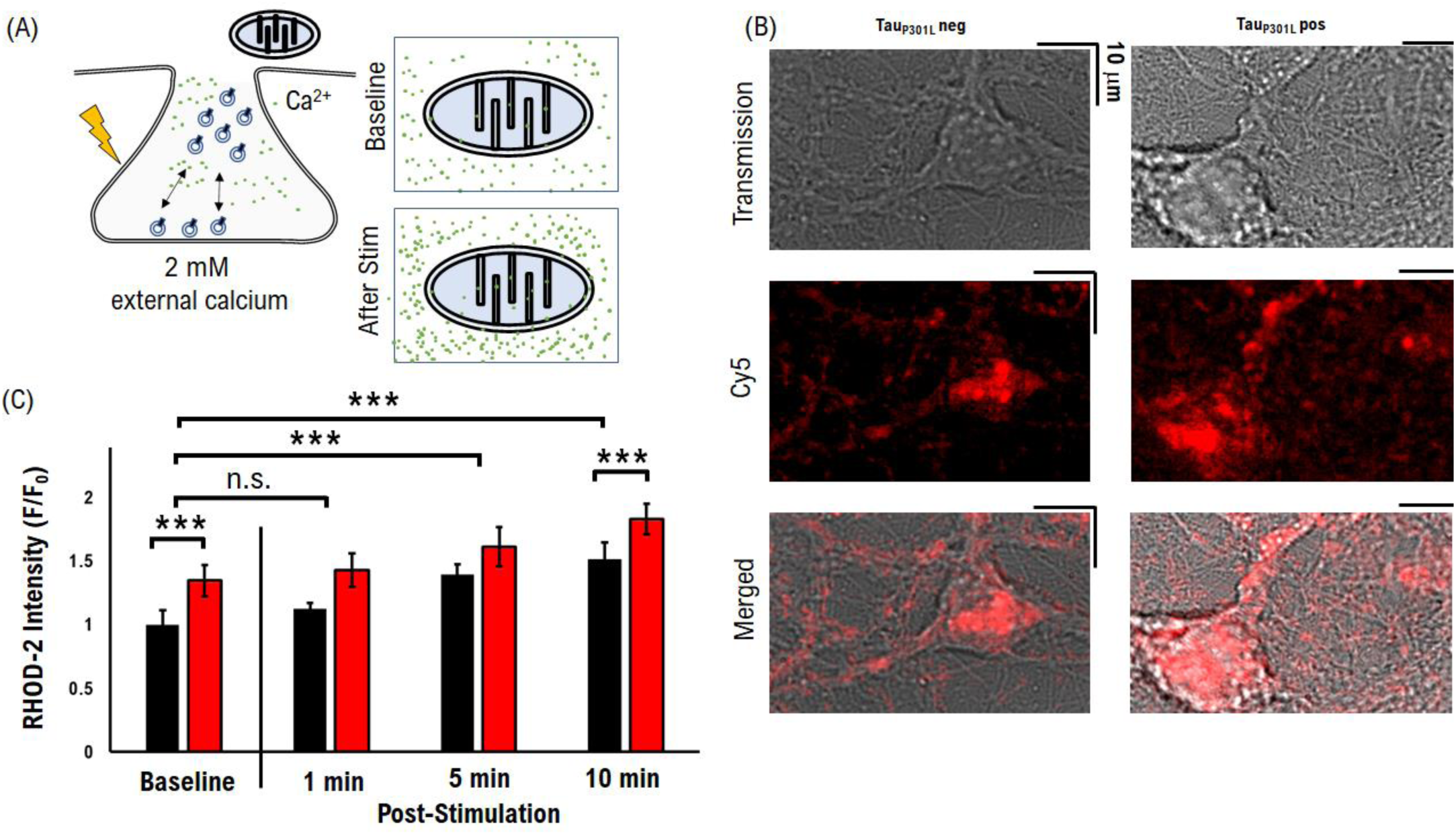
TauP301L pos mitochondria absorb more calcium during high activity. (A) Model of increased calcium absorption measured via RHOD-2 (B) Example images of measured RHOD-2 intensity for tau_P301L_ neg (left column) and tau_P301L_ pos (right column) cultures. RHOD-2 puncta co-localize with neuronal processes observed with transmission imaging. (C) Relative RHOD2 integrated intensity of tau_P301L_ neg (Black) and tau_P301L_ pos (Red) mitochondria. Tau_P301L_ neg: N >4000, 3 samples, 1 litter. Tau_P301L_ pos: N >4000, 3 samples, 1 litter. *** = P < 0.001;

We observed that tau_P301L_ pos mitochondria exhibited slightly elevated RHOD-2 intensity (**Red**, **Fig. 7C**), at baseline compared to tau_P301L_ neg mitochondria (**Black**, **Fig. 7C**), indicating elevated calcium-absorption (P<0.001). After a bout of high stimulation, both tau_P301L_ pos and tau_P301L_ neg mitochondria calcium concentration increases at proportionally the same rate (**Fig. 7C**). At 10 min after stimulation tau_P301L_ pos mitochondria have slightly elevated amount of absorbed calcium (1.77 +/- 0.2) compared to tau_P301L_ neg mitochondria (1.46 +/- 0.1). Further, we observed that after a bout of high-activity, in the absence of external calcium, rescued the time-dependent change in membrane potential (Fig. S2).

These combined results suggest that differences in adsorbed calcium is not the main contributing factor to observed differences in mitochondria membrane potential after high-activity (**Fig. 5D**).

### Reduced glutamate Concentration Per Vesicle Rescues Activity-dependent Changes in Tau_P301L_ pos Mitochondria Membrane Potential

To test the role of glutamate recycling on mitochondria function, we reduced ATP-dependent glutamate refilling into synaptic vesicles via VGlut1. Vesicle glutamate refilling represents a large portion of energy demand during recycling.^11, 60^ If vesicle glutamate concentration increases due to the increase of VGlut1 transporters per vesicle, then the amount of ATP-dependent demand would also increase. To test the role of ATP-dependent vesicle refilling, we used an established competitive VGlut1 ATP-inhibitor Trypan Blue (**Fig. 8 A**).^60^ Trypan Blue (TB) binds to VGlut1 and inhibits ATP-dependent glutamate refilling, thus reducing ATP demand. If reduced glutamate-dependent ATP demand reduces activity-dependent mitochondria membrane potential, then the observed difference in tau_P301L_ pos mitochondria membrane potential is due to differences in ATP demand.

**Fig 8:**
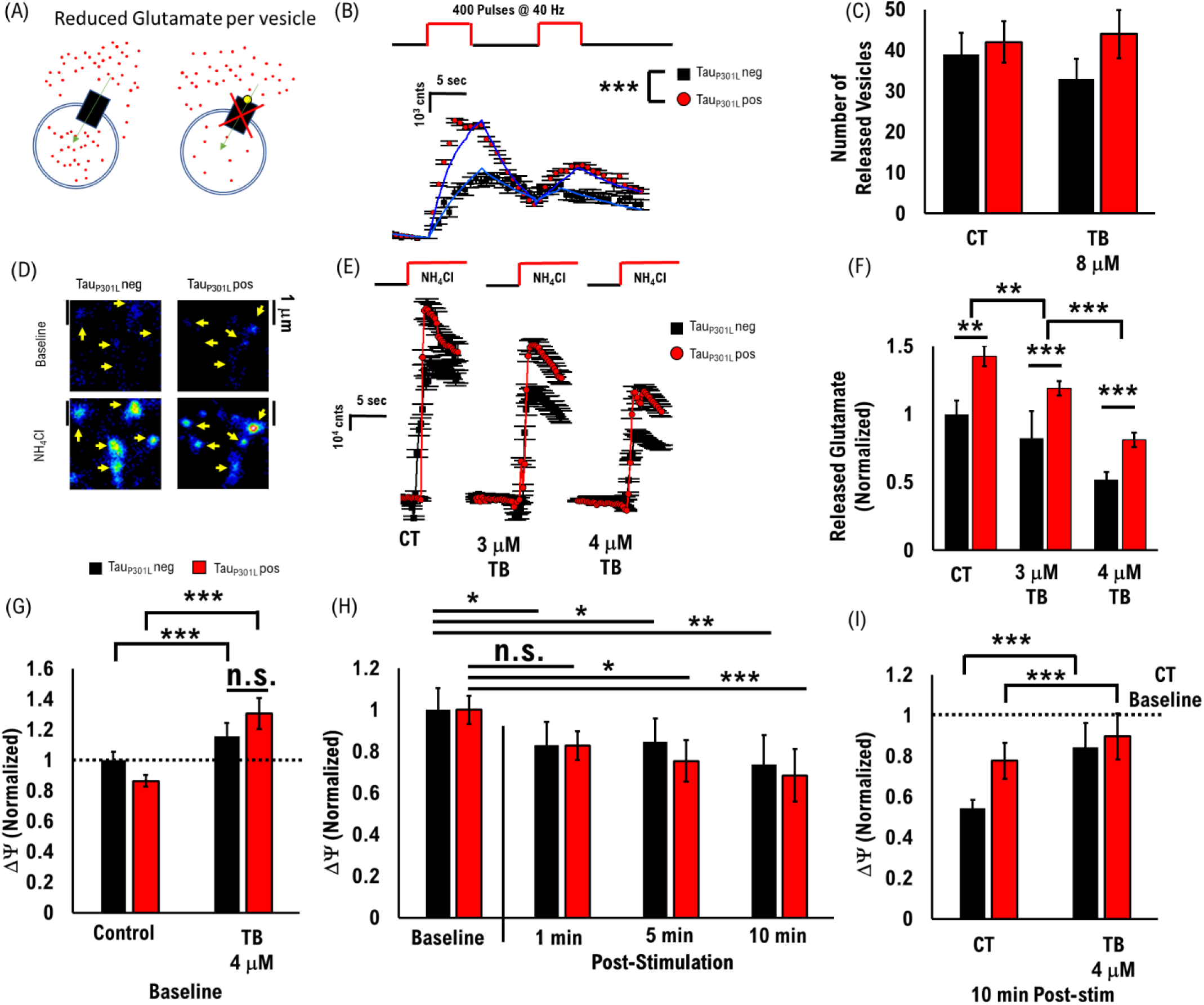
Activity-dependent changes in Mitochondria Membrane potential for 0mM external Calcium. (A) Cartoon model of TB inhibition of VGLUT1-dependent glutamate transport (B) VGlut1-pHluorin intensity during two bouts of 40 Hz stimulation in the presence of 4 μM TB for tau_P301L_ pos (Red) and tau_P301L_ neg (Black). Computational model fits (Blue lines) reproduce experimental results. (C) Number of released vesicles obtained from computational model fits for each condition. Vesicles are from two bouts of 40Hz stimulation data in (B). (D) Example images of iGluSnFr fluorescence before and during exposure to NH4Cl. (E) Time-dependent iGluSnFr fluorescence during exposure to NH4Cl. (F) Peak iGluSnFr intensity during exposure to NH4Cl in the presence of 3 μM or 4 μM TB for tau_P301L_ pos (Red) and tau_P301L_ neg (Black). (G) Relative baseline mitochondria membrane potential in the presence of 4 μM TB relative to 2mM external Ca^2+^ (H) Relative change in mitochondria membrane potential in the presence of 4 μM TB as a function of time after a bout of high-activity (I) Relative change in mitochondria membrane potential 10 min after activity in the presence of 4 μM TB relative to 2mM external Ca^2+^ Tau_P301L_ neg: VGlut1-pHluorin data:; iGluSnFr data: N = ;JC-1 data: N >5000, 5 samples, 2 litters. Tau_P301L_ pos: VGlut1-pHluorin data: ; iGluSnFr data: N = ;JC-1 data: N >6000, 5 samples, 2 litters. *= P < 0.06; ** = P<0.01; *** = P < 0.001; pHluorin statistical comparisons are Mann-Whitney U test. JC-1 comparisons within rTg conditions are two-tailed KS-test of cumulative distributions. JC-1 comparisons across rTg conditions are pair-wise two-tailed t-Test.

We first tested the number of vesicles released and the amount of glutamate released during activity to support the hypothesis that TB does not alter vesicle release mechanics but does reduce glutamate concentration per vesicle. We used a high concentration (0.8 μM TB), compared to previous in vitro studies,^60^ to show that VGlut1-pHluorin intensity was not differentially affected for tau_P301L_ neg (Black, **Fig. 8 B**) or tau_P301L_ pos (Red, **Fig. 8 B**) synapses compared to control VGlut1-pHluorin intensity (**Fig. 5 B**). We then measured the amount of glutamate released as a function of increasing TB concentration using the established glutamate reporter iGluSnFr (**Fig. 8 D**), in a range consistent with in vitro studies,^60^ using our previously established approach.^8^ Here, we found a reduction in the amount of released glutamate relative to controls for both tau_P301L_ neg (Black, **Fig. 8 E**) and tau_P301L_ pos (Red, **Fig. 8 E**) synapses. The reduction was proportional to the increase in TB concentration and was the same rate for both tau_P301L_ pos and tau_P301L_ neg neurons (**Fig. 8F**). Further, the total amount of glutamate release was approximately half in the presence of 0.4 μM TB compared to baseline (**Fig. 8F**), which is consistent with previous in vitro studies.^60^ These results show that exposure to TB reduces the amount of glutamate released without altering the mechanics of vesicle release.

We next measured the effects of reduced glutamate refilling per vesicle on mitochondria membrane potential at baseline and after a bout of high activity. We first observed that baseline membrane potential was higher for both tau_P301L_ pos and tau_P301L_ neg mitochondria in the presence of 4 μM TB compared to control (**Fig. 8 G**). However, the change in membrane potential as a function of time after high activity showed that both tau_P301L_ neg (Black, **Fig. 8 H**) and tau_P301L_ pos (Red, **Fig. 8 H**) mitochondria followed the same time-dependent reduction as observed under control conditions (**Fig. 5E**). Further, both tau_P301L_ neg and tau_P301L_ pos membrane potential were the same 10 min after activity (**Fig. 8I**) in the presence of 4 μM TB compared control baseline without TB.

Taken together, these results suggest that mitochondria function is dynamically changing in order to support increased glutamate release and recycling in tau_P301L_ pos neurons during hyperexcitable states.

### Hypothesized Model of Dynamic Mitochondria Function in P301L Neurons

We now hypothesize a model of mitochondria membrane potential changes under hyper-excitable conditions based on combined La-SEM and membrane potential results (**Fig. 9**). Our model proposes that ATP-demand represents the dominant influence on mitochondria structural and functional changes observed in tau_P301L_ pos neurons prior to the observable reduction in presynaptic function. Our model also proposes that increased ATP-generating cristae centers allow tau_P301L_ pos mitochondria to maintain a higher membrane potential during periods of extended activity due to their ability to support higher ATP demand (**Fig. 9C**). Further, late-stage changes in mitochondria calcium absorption are not a significant contributing factor to mitochondria function in tau_P301L_ pos cultured neurons.

**Fig 9:**
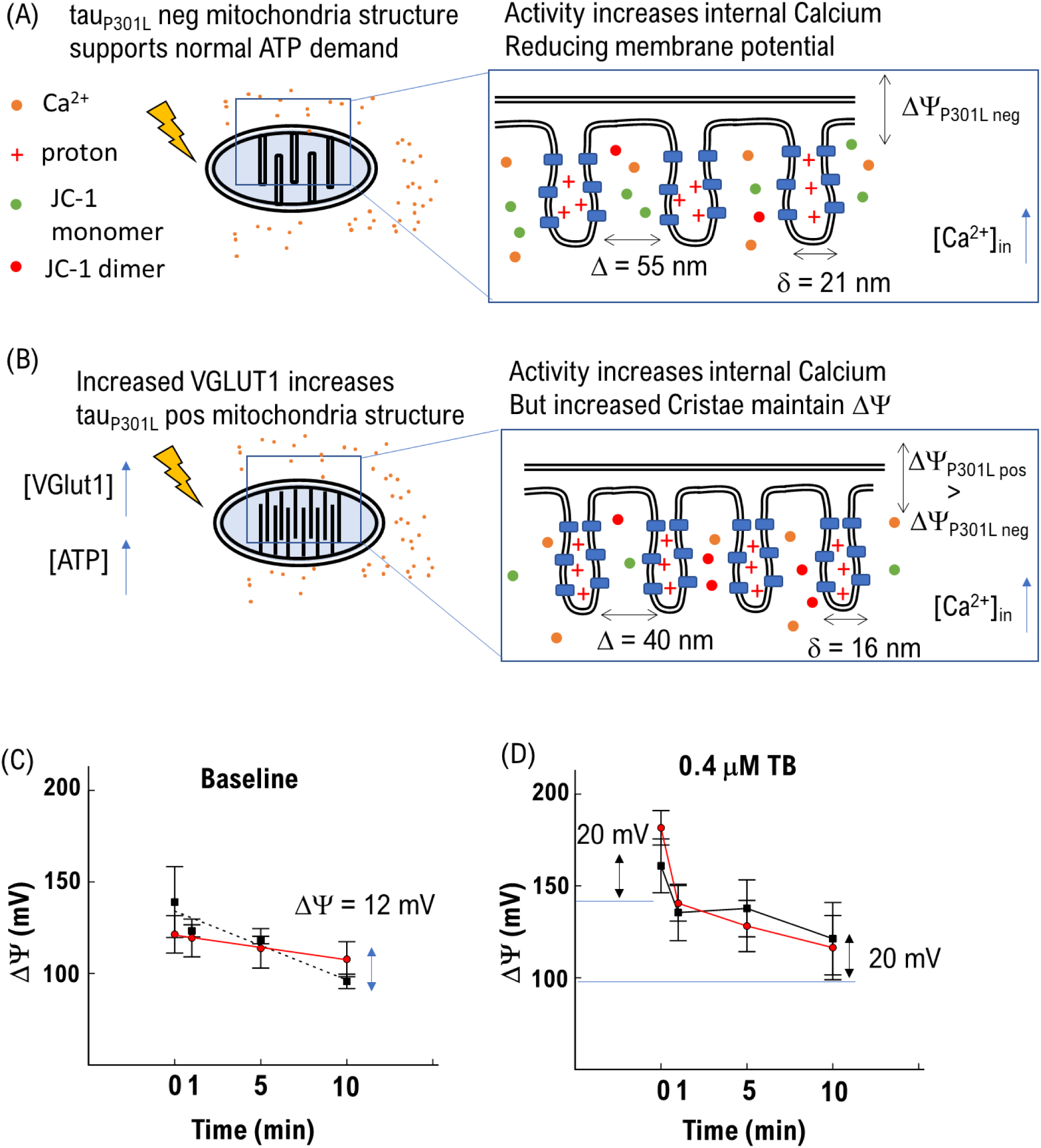
TauP301L pos mitochondria absorb more calcium during high activity. (A) Model of Mitochondria function under for tau_P301L_ neg neurons during activity based on observed La-SEM structure (B) Model of Mitochondria function under for tau_P301L_ pos neurons during activity based on observed La-SEM structure (C) Change in membrane potential for tau_P301L_ neg (Black) and tau_P301L_ pos (Red) mitochondria under control conditions. (D) Change in membrane potential for tau_P301L_ neg (Black) and tau_P301L_ pos (Red) mitochondria in the presence of 0.4 μM TB.

To support this model, we first compare differences in ATP generating cristae centers. Under normal conditions, mitochondrial support ∼13 ATP generation and transport centers per micron length via a ∼55 nm cristae distance and 21 nm cristae thickness (**Fig. 9A**). Increased VGlut1 and glutamate recycling demand during homeostasis in tau_P301L_ pos neurons increases ATP generation demand in mitochondria resulting in an increase in ATP generating cristae to ∼16 per micron length, via a reduction in cristae distance to 45 nm and thickness of 16 nm (**Fig. 9B**). Thus, tau_P301L_ pos mitochondria have 20-30% more ATP generating cristae centers compared to tau_P301L_ neg mitochondria.

We also support this model by comparing changes in membrane potential under controlled conditions after high-activity. Adsorbed calcium alone is sufficient to explain changes in membrane potential in tau_P301L_ neg mitochondria, following a Nearnst equilibrium potential model and a total adsorbed calcium amount of ∼500 nM. However, calcium adsorption alone is insufficient to explain the observed increase in tau_P301L_ pos mitochondria membrane potential, because increased membrane potential occurs even while calcium adsorption is increased (**Fig. 7**). Alternatively, reduced ATP demand, mediated by reducing glutamate uptake via VGlut1 inhibition, results in an increase in mitochondria membrane potential during resting state and after high-activity equally for both tau_P301L_ neg and pos mitochondria (**Fig. 9D**). The increase in membrane potential is caused by the rise in internal ion concentration relative to the cytosol due to reduced ATP production. However, calcium absorption still dominates mitochondria membrane potential after activity, but tau_P301L_ pos and tau_P301L_ neg mitochondria have the same membrane potential due to reduced ATP demand (**Fig. 9D**).

## Conclusion

We observed a significant difference in tau_P301L_ pos mitochondria morphology showing longer (length) but more narrow (width) relative to tau_P301L_ neg mitochondria (**Fig 1**). This difference in length-to-width change corresponded with an internal structural change where tau_P301L_ pos mitochondria narrower cristae that are more closely packed together (**Fig. 2**). We observed that the change in morphology and structure corresponded with an increase in the density of membrane material, but no significant change in the amount of adsorbed calcium granules (**Fig. 3**). We next showed that the baseline structural change also corresponded with a change in the resting mitochondria membrane potential (**Fig. 4**). This change in resting membrane potential was not due to changes in baseline presynaptic vesicle release mechanics (**Fig. 4**). We hypothesized that the difference in mitochondria membrane potential may be due to differences observed in the amount of glutamate concentration per vesicle (**Fig. 4**). We tested this hypothesis by measuring differences membrane potential as a function of time after a bout of low-activity (one train of 100 pulses at 10 Hz), or a bout of high-activity (two trains of 400 pulses at 40 Hz). We observed that mitochondrial membrane potential initially decreased with time for both low and high activity levels (**Fig. 5**). The membrane potential returned to baseline 10 min after low-activity (**Fig. 5**). In contrast, the membrane potential remained lower 10 min after the high activity (**Fig. 5**). We measured calcium-absorption after high-activity using RHOD-2 indicator and showed that membrane potential differences observed in tau_P301L_ pos mitochondria are not dependent on calcium absorption (**Fig. 7**). We then rescued the difference in membrane potential between tau_P301L_ neg and tau_P301L_ pos mitochondria by inhibiting synaptic vesicle glutamate loading via the competitive VGlut1 agonist Trypan blue and observed the same baseline membrane potential and time-dependent membrane potential after high-activity (**Fig 8**). Finally, we propose a hypothesized model that differences in mitochondria membrane potential for tau_P301L_ pos neurons is due to the increased cristae density observed in tau_P301L_ pos mitochondria that produce more ATP to support increased energy demands.

## Discussion

### Mitochondria Morphology and Membrane Density in Early Tauopathy

It is well established that mitochondria rapidly change their morphology and structure to support changing energy demands within cells.^41, 61^ However, it is also well established that mitochondria have altered morphology and function during tauopathy progression,^17, 24, 62^ which has been hypothesized to be directly mediated by altered forms of tau. Thus, it is important to separate mitochondria changes that occur due to changes in energy demands versus direct influence by tau.

Our observations of altered mitochondria morphology and function in the presence of P301L (Fig. 1-5) follows the established understanding that mitochondria are altered during tauopathy, and have increased Ca^2+^ absorption in the presence of P301L tau (Fig. 8). But our observation of rescued mitochondria function, by reducing VGlut1 mediated ATP demand (Fig. 7), also supports the established dynamics of mitochondria in normal function. This rescued effect suggests that, at least in early stages of disease progression, altered mitochondrial function and Ca^2+^ absorption are secondary affects due to increased energy demand in P301L tau disease.

These results lead to several pathways that warrant future study. First, indirect early-stage alterations in mitochondrial morphology and function may lead to faster mitochondrial degradation and be a contributing factor to later stage reduction in synaptic transmission, in parallel with direct tau-mediated changes in synaptic function, that exacerbate neurodegeneration (which we discuss below). Second, increased mitochondrial function supports increased synaptic transmission that supports transsynaptic spread of tau, and thus indirectly mediate disease progression (discussed below). Third, increased mitochondria Ca^2+^ absorption and function require more demand from neurons to maintain homeostasis resulting in faster breakdown of neuronal function indirectly, and thus contributing to neurodegeneration.

### Mitochondria Demand Is a Parallel Pathway to Neurodegeneration

It is important to consider the increased mitochondria function observed in this present work in the context of normal aging in order to better understand how early changes in mitochondria can lead to neurodegeneration. Mitochondria ATP synthesis utilizes a large combination of protein function and interactions.^27^ Proteins regularly break down with use and time,^28, 29^ and must be regularly replaced.^29^ Fusion and fission have been shown to be an essential part of mitochondrial replenishment in order to maintain proper function.^63^ Breakdown in mitochondria function through protein degradation has been shown to lead to mitochondrial degeneration and ultimately neural degeneration in multiple different pathways.^30^ These established results show that increasing mitochondrial use is a potential pathway to neurodegeneration.

In the context of mitochondrial aging, our observed increase in mitochondrial demand prior to observed changes in presynaptic function supports the hypothesis that increased glutamate recycling (release, clearance, refilling) leads to neurodegeneration. The increased glutamate recycling process, via tau-mediated increased VGlut1 expression, increases demand on mitochondria to support increased ATP use. This increase in mitochondria demand leads to altered mitochondria morphology and function, which in the short-term is a net positive because it supports increased demand. However, increased mitochondrial use is a long-term negative process as it results in faster protein degradation. Replacing mitochondria that age faster is thus an extra burden on neurons that over time may not be capable of maintaining this rate of replenishment. Thus, the ultimate result of this pathway means that mitochondria aging occurs faster and is a parallel pathway, with tau-mediated changes in presynaptic transmission, that exacerbates neuronal dysfunction during disease progression.

### VGlut1 as a target to reduce or arrest neurodegeneration in P301L tauopathy

As we have shown, tau-mediated increase in VGlut1 transporters per vesicle results a significant increase in overall demand from mitochondria (Fig. 5). We have also shown that inhibiting VGlut1 activity, via the competitive blocker Trypan Blue, rescues effects from tau-mediate mitochondria changes (Fig. 7). These results suggests that reducing glutamate demand is a potential therapeutic target to reduce or arrest disease progression. This pathway has two potential advantages over other approaches:

First, increased VGlut1 expression by P301L has recently been shown to occur by tau directly influencing gene expression, via epigenetic factors TRIM28 and HDAC1.^9^ To prevent or inhibit tau mediate changes in gene expression machinery therapeutically would require directly targeting these nuclear compartment factors. However, targeting gene expression in the nuclear compartment could potentially have off-target affects resulting in altered gene expression of other essential proteins for neuronal function, and will take significant work to achieve. Further, inhibiting altering expressing in patients that already express increased VGlut1 may not reduce the effects currently ongoing, such as hyperexcitability. As a parallel approach, targeting the downstream effects of increased glutamate release via VGlut1 could have the benefit of reducing ongoing disease symptoms and arrest the progression of the disease.

Second, activity mediated trans-synaptic spread of tau is now an established pathway of tau spread throughout the brain,^31, 64^ which is exacerbated by the hyper-excitable state resulting in increased synaptic activity. Reducing glutamate release by targeting VGlut1 directly represents a pathway to reduce trans-synaptic tau spread. If hyperexcitability is mediated by increased glutamate release, then reducing glutamate would prevent hyperexcitability from occurring. Inhibiting hyperexcitability, would then reduce the amount of network activity resulting in reduced tau release. While this approach would not completely arrest activity-mediated release of tau, it would represent a pathway to reduce its spread and slow disease progression.

## Acknowledgements

We acknowledge the assistance of Dr. Sanja Sviben and Greg Strout at the Washington University Center for Cellular Imaging (WUCCI) in EM studies, which is supported by Washington University School of Medicine, The Children’s Discovery Institute of Washington University and St. Louis Children’s Hospital (CDI-CORE-2015-505 and CDI-CORE-2019-813) and the Foundation for Barnes-Jewish Hospital (3770 and 4642). We would like to thank Washington University Hope Center Viral Vector Core for the production of the Glut1-pHluorin vector used in this study.

## Author contributions

MG and MR performed the experimental design and analysis. RC managed the mouse colony breeding, and sample preparation. MG, RC, MH, DG, and JP performed the experiments. MG, MH, DG and MR analyzed the experimental results and wrote the manuscript. MG performed the computational analysis. All authors contributed to the article and approved the submitted version.

## Supplementary Material

**Fig S1:**
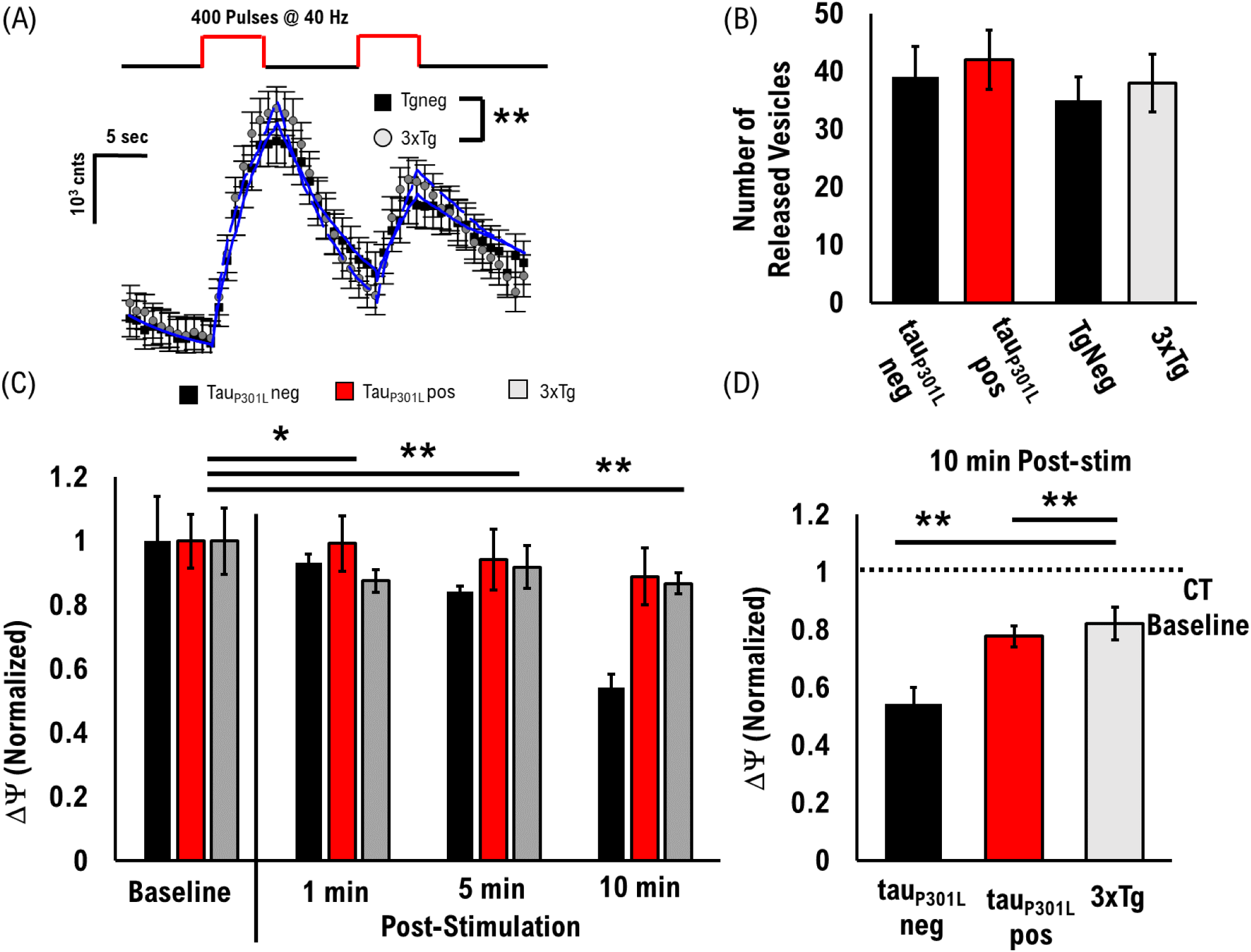
3xTg neurons exhibit similar VGLUT1 and Mitochondria Changes as Tau_P301L_ pos neurons. (A) VGlut1-pHluorin intensity during two bouts of 40 Hz stimulation for 3xTg neurons (Grey) and Tgneg (Black). Computational model fits (Blue lines) reproduce experimental results. (B) Number of released vesicles obtained from computational model fits for each condition. Vesicles are from two bouts of 40Hz stimulation data in (A). (C) Relative change in mitochondria membrane potential as a function of time after a bout of high-activity (D) Relative change in mitochondria membrane potential 10 min after activity normalized to tau_P301L_ neg baseline VGlut1-pHluorin: TgNeg, N 150, 1 samples, 1 litter; 3xTg N 416, 1 samples, 1 litter; ** = P<0.01; Mann-Whitney U test JC-1: N > 3500; 3 samples, 1 litter; ** = P<0.01; two-tailed KS-test

To determine if changes mediated by the P301L tau gene alone are sufficient to alter membrane potential differences, we utilized the 3xTg mouse model which includes the P301L tau mutation, a PS1_M146V_ mutation, and an APP_Swe_ mutation.^65^ Mice with these three mutations combined exhibit altered glutamate release as well as other late-stage cognitive declines.^65^ These mice develop Aβ plaques and intraneuronal Aβ immunoreactivity during late-stages of the disease progression. However, it is not clear if the other mutations also influence mitochondria function. To test this possibility, we first confirmed that VGlut1 is increased under the same cell culture conditions and observed an increase in overall VGlut1-pHluorin intensity consistent with the tau_P301L_ pos neurons (Fig. S1 A), but not a significant difference in the number of release vesicles (Fig. S1 B). We note that we could not compare absolute values at baseline JC-1 intensities as the samples were taken under different experimental conditions from the tau_P301L_ cultures, but we did compare the relative change in average ratio of 3xTg JC-1 to relative change in tau_P301L_ cultures. We observed a similar reduction in baseline JC-1 intensity ratio for 3xTg (Fig. S1 C), although less significant that observed for tau_P301L_ pos neurons. Importantly, we observed the same time-dependent change in membrane potential after high-activity (Grey Fig. S1 D), as observed with tau_P301L_ pos neurons (Red Fig. S1 D). This difference resulted in the same elevated membrane potential at 10 min after high activity as observed for tau_P301L_ pos neurons (Fig. S1 E). These results suggest that the PS1_M146V_ mutation, and an APP_Swe_ mutations do not significantly affect mitochondrial membrane potential during early stages of disease progression.

### Difference in membrane potential for tau_P301L_ pos mitochondria is driven by Calcium adsorption

**Fig S2:**
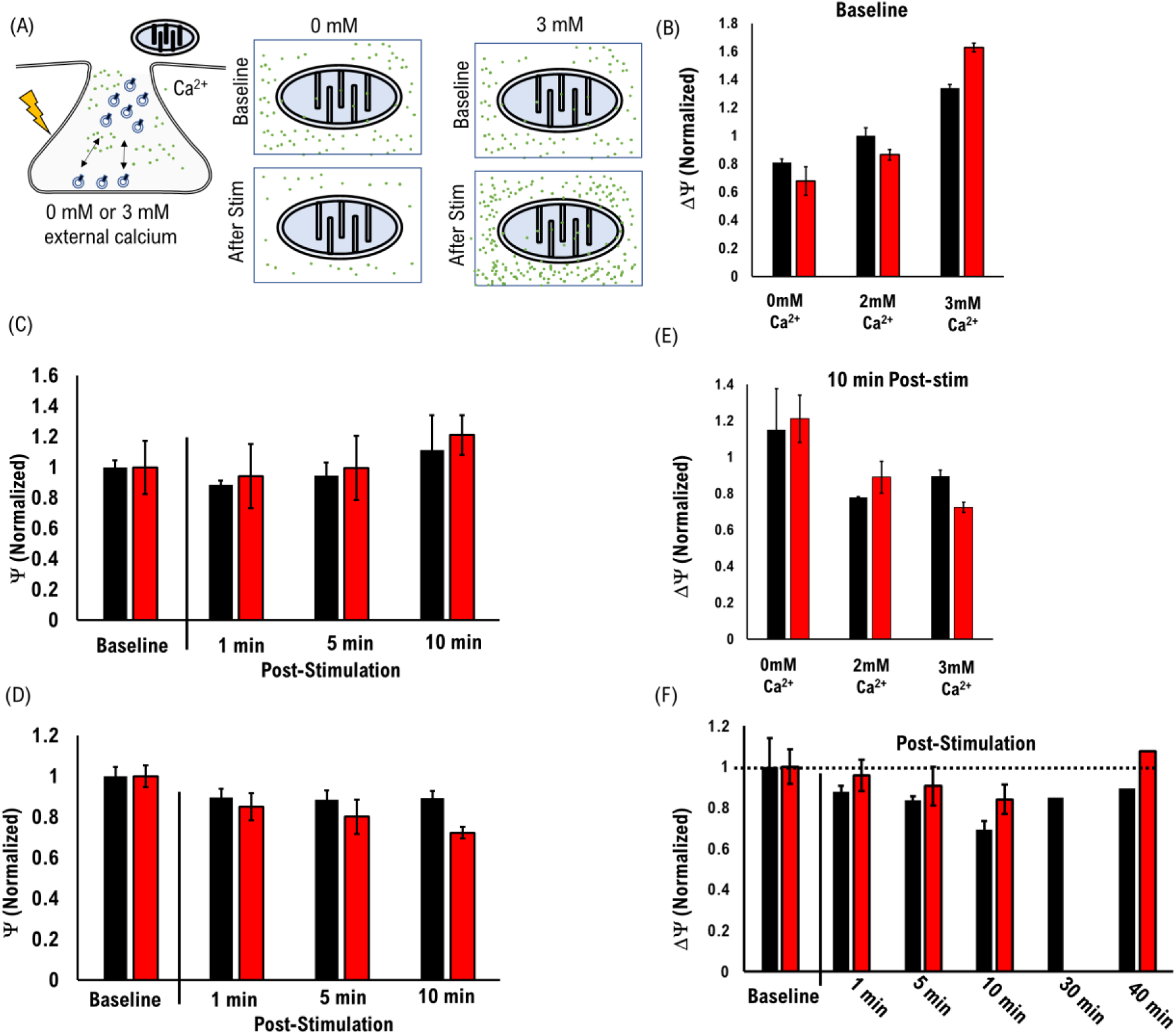
Activity-dependent changes in Mitochondria Membrane potential for 0mM external Calcium. (A) Cartoon model of changing internal calcium concentration affect of mitochondria with 0mM or 3mM external calcium (B) Relative baseline mitochondria membrane potential in the presence of 0mM, 2mM, 3mM external Ca^2+^ (C) Time-dependent change in mitochondria membrane potential with 0mM external calcium, after a bout of high-activity relative to baseline. (D) Time-dependent change in mitochondria membrane potential with 3mM external calcium, after a bout of high-activity relative to baseline. (E) Relative mitochondria membrane potential 10 min after a bout of high-activity in the presence of 0mM, 2mM, 3mM external Ca^2+^ and normalized by resting potential of each group. (F) Time-dependent change in mitochondria membrane potential with 2mM external calcium, after a bout of high-activity relative to baseline, showing the timescale of return to baseline. 0mM: tau_P301L_ neg: N>1000 mitochondria per sample from 2 plates and 1 litter. 0mM: tau_P301L_ pos: N>1000 mitochondria per sample from 2 plates and 1 litter. 3mM: tau_P301L_ neg: N>1000 mitochondria per sample from 2 plates and 1 litter. 3mM: tau_P301L_ pos: N>1000 mitochondria per sample from 2 plates and 1 litter.

To test the hypothesis that time-dependent change in membrane potential is influenced by external calcium, we measured mitochondria membrane potential as a function of time after a bout of high-activity, but either in the absence of external calcium (**Fig. S2 A**), or with greater external calcium. By changing external calcium concentration and not internal calcium concentration, internal calcium concentration will change after activity in order to equilibrate with external calcium resulting in a changing amount of mitochondria absorbed calcium with time. If changes in mitochondria membrane potential are calcium-dependent, then this approach should show changes in membrane potential relative to 2mM external calcium.

We first measured changes in mitochondria membrane potential at resting to determine if mitochondria were adversely affected by external calcium concentration (**Fig. S2 B**). We observed a clear increase in resting membrane potential with increasing external calcium concentration for both tau_P301L_ neg (**Black, Fig. S2 B**) and tau_P301L_ pos (**Red, Fig. S2 B**) mitochondria. This effect was the same for both tau_P301L_ neg or tau_P301L_ pos cultures showing that this does not adversely affect tau_P301L_ pos.

We then quantified time-dependent changes in mitochondria membrane potential after a bout of high-activity the absence of external calcium concentrations. For 0mM external calcium, both the tau_P301L_ neg (**Black, Fig. S2 C**) and tau_P301L_ pos (**Red, Fig. S2 C**) mitochondria show an initial reduction in membrane potential 1 min after stimulation, followed by a return to the same membrane potential 5 min after stimulation. This would be consistent with recovery after the initial release of vesicles observed even in the absence of external calcium (Supplementary Material). However, unlike in the presence of 2mM external calcium, both tau_P301L_ neg and tau_P301L_ pos mitochondria exhibit increasing membrane potential as a function of time relative to baseline in the absence of external calcium. As a result, 10 min after activity, both tau_P301L_ neg (**Black, Fig. S2 E**) and tau_P301L_ pos (**Red, Fig. S2 E**) mitochondria have greater membrane potential relative to 10 min after activity in the presence of 2mM external calcium. Moreover, both tau_P301L_ pos mitochondria membrane potential and tau_P301L_ neg membrane potential are the same relative change in Δψ, which shows tau_P301L_ pos mitochondria are not adversely affected by changes in calcium concentration.

We next quantified the time-dependent change in mitochondria membrane potential after a bout of high-activity the presence of 3mM external calcium concentrations. Both the tau_P301L_ neg (**Black, Fig. S2 D**) and tau_P301L_ pos (**Red, Fig. S2 D**) mitochondria show a consistent reduction in membrane potential with time after stimulation, and both are consistent with the observed decrease in the presence of 2mM external calcium. Further, 10 min after activity, both tau_P301L_ neg (**Black, Fig. S2 E**) and tau_P301L_ pos (**Red, Fig. S2 E**) mitochondria have similar membrane potential relative to 2mM external calcium. However, unlike 2mM external calcium, tau_P301L_ neg mitochondria membrane potential is slightly larger than tau_P301L_ pos membrane potential at 3 mM external calcium, which suggests that tau_P301L_ pos mitochondria are adversely affected by changes in calcium concentration.

### Astrocyte Glutamate recycling does not modulate Tau_P301L_ pos mitochondria membrane potential

**Fig S3:**
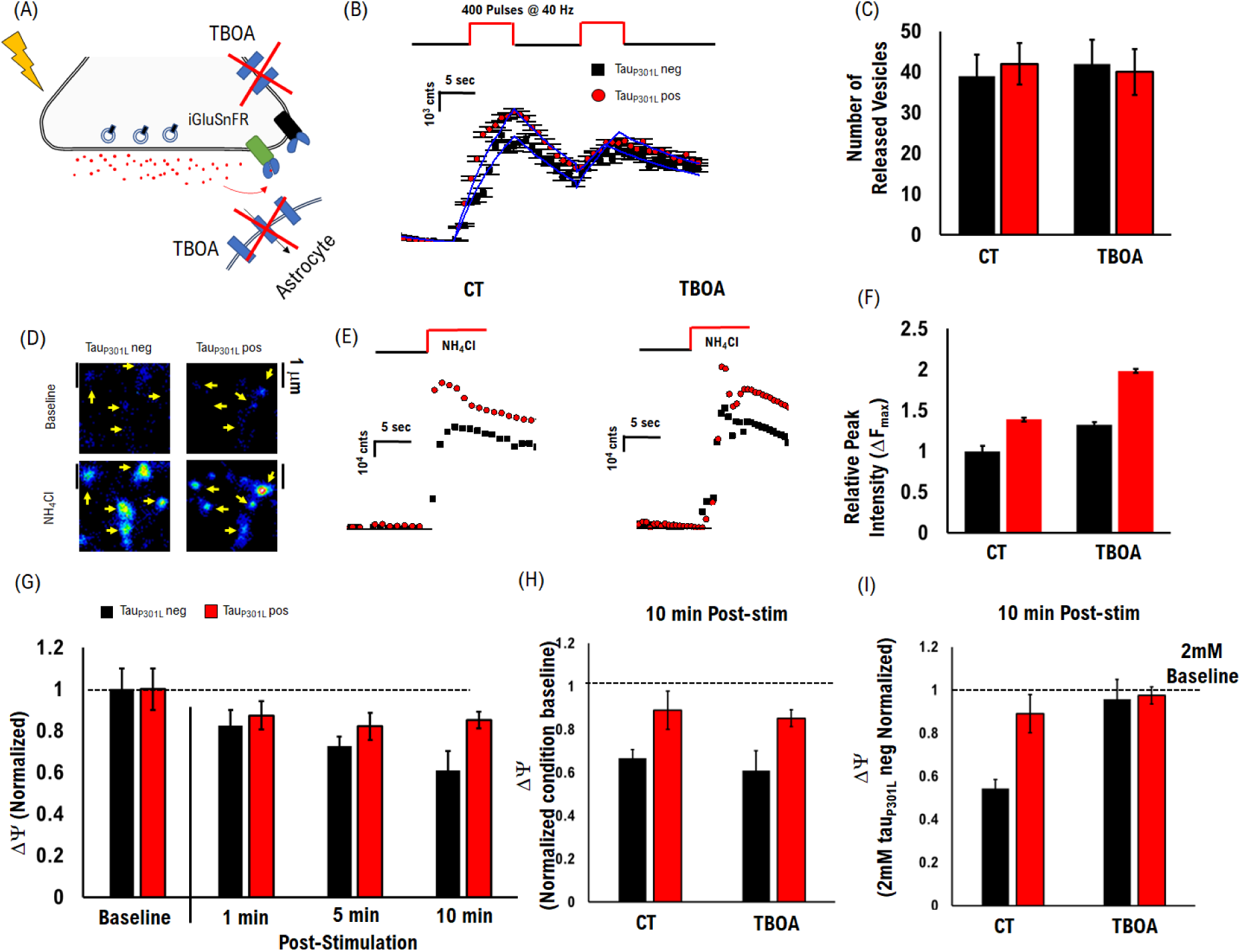
Glutamate uptake Affects Membrane Potential Scale but not Activity-dependent Ca^2+^ absorption. (A) Cartoon model of TBOA inhibition of glutamate reuptake and resulting iGluSnFr receptor response (B) VGlut1-pHluorin intensity during two bouts of 40 Hz stimulation in the presence of TBOA for tau_P301L_ pos (Red) and tau_P301L_ neg (Black). Computational model fits (Blue lines) reproduce experimental results. (C) Number of released vesicles obtained from computational model fits for each condition. Vesicles are from two bouts of 40Hz stimulation data in (B). (D) Example iGluSnFr fluorescence images before and after exposure to NH4Cl. (E) iGluSnFr fluorescence intensity as a function of time after exposure to NH4Cl for control conditions (left) and under exposure to the glutamate reuptake inhibitor TBOA (right). (F) Quantified peak iGluSnFr fluorescence intensity differences for control and TBOA exposed cultures. Peak intensity increases in the presence of TBOA indicating slower glutamate clearance after activity. (G) Relative JC-1 measured mitochondria membrane potential under High Stimulation shows a significant and sustained decrease for Tau_P310L_ neg (Black) neurons after stimulation, but a less significant decrease for Tau_P310L_ pos (Red). (H) Relative JC-1 measured mitochondria membrane potential 10 min after high stimulation, normalized to each conditions baseline, shows the same time-dependent relative reduction for both tau_P310L_ neg (Black) mitochondria and tau_P310L_ pos (Red), compared to control. (I) Relative JC-1 measured mitochondria membrane potential 10 min after high stimulation normalized to tau_P301L_ neg control shows the same membrane potential as observed for the 0.4 μM TB inhibition of VGLUT1 (Fig. 7 I). Tau_P301L_ neg: VGlut1-pHluorin data: N = 684, 2 samples, 1 litter; iGluSnFr data: N = 169, 2 samples, 1 litter;JC-1 data: N >3000, 3 samples, 1 litters. Tau_P301L_ pos: VGlut1-pHluorin data: N = 1085, 2 samples, 1 litter; iGluSnFr data: N = 150, 2 samples, 1 litter;JC-1 data: N >4000, 3 samples, 1 litters. ** = P<0.01; *** = P < 0.001; pHluorin/iGluSnFr statistical comparisons are Mann-Whitney U test.

To determine if our observed mitochondria changes were a consequence of glutamate uptake of extracellular glutamate. Glutamate clearance is the later stage of the recycling process after release. If more glutamate is released in tauP301L pos neurons compared to tauP301L neg neurons, then the ATP-dependent reuptake of extracellular glutamate could potentially increase demand from mitochondrial ATP production. This increased demand would then be a contributing factor to the activity-dependent reduction in mitochondria membrane potential observed here (Fig. 5). To determine the role of glutamate re-uptake on mitochondria membrane potential via ATP demand, we used the established acute blocker (TBOA, Fig. S3 A), which inhibits excitatory amino acid transporter-1 (EAAT-1) from clearing extracellularly released glutamate.^66–68^

To confirm that TBOA affects the re-uptake pathway without inhibiting glutamate release, we first determined if the TBOA inhibitor reduced either the number of vesicles released or the amount of glutamate released prior to re-uptake. We quantified the amount of released vesicles under the high stimulation condition by measuring VGlut1-pHluorin intensity in the presence of TBOA (Fig. S3 B). We observed the same increase in intensity for tauP301L pos neurons compared to tau_P301L_ neg neurons showing that TBOA did not adversely affect the relative difference in vesicle release. The number of released vesicles was unchanged in the presence of TBOA compared to control conditions (Fig. S3 C).

We next determined if TBOA altered the amount of glutamate released extracellularly using our previously established approach using the external glutamate reporter iGluSnFr (Fig. S3 D).^8^ We compared peak iGluSnFr intensity to determine any changes in the amount of extracellular glutamate in the absence (CT, Fig. S3 E,F) or presence (TBOA, Fig. S3 E,F) of TBOA. We found that iGluSnFr peak intensity increased in the presence of TBOA by ∼32 tau_P301L_ neg neurons compared to control, and by 50% for tau_P301L_ pos compared to control (Fig. S3 E,F). This result is consistent with the mechanism of TBOA: TBOA reduces glutamate clearance which would increase the local concentration of external glutamate in the synaptic cleft. These results show that TBOA does not alter vesicle release mechanics or the amount of glutamate released, but does reduce glutamate clearance by 30-50%.

To determine the effects of reduced glutamate re-uptake on mitochondria membrane potential, we followed the same high-stimulation activity-dependent JC-1 membrane potential protocol used for control, but in the presence of TBOA (Fig. S3 G-I). We observed the same relative reduction in membrane potential after the high stimulation protocol, for both the tau_P301L_ pos and tau_P301L_ neg mitochondria compared to control (Fig. S3 G). The resulting membrane potential 10 min after the high stimulation protocol, normalized to each conditions baseline, was consistent with the relative membrane potential under control conditions (Fig. S3 H). However, when normalizing the mitochondria membrane potential to the tau_P301L_ neg control baseline, we observed the same mitochondria membrane potential for both tau_P301L_ neg and tau_P301L_ pos observed when we inhibited VGlut1 using 0.4 μM TB (Fig. S3 I).

Taken together these results suggest that glutamate reuptake energy requirements do affect local mitochondria baseline membrane potential, but not the time-dependent potential after a bout of high-activity. We note, however, that there are a large number of processes in the pathway involved in glutamate uptake that can have compounding effects on mitochondria membrane potential.

Computational Model Parameters Table:

**Table S1:**
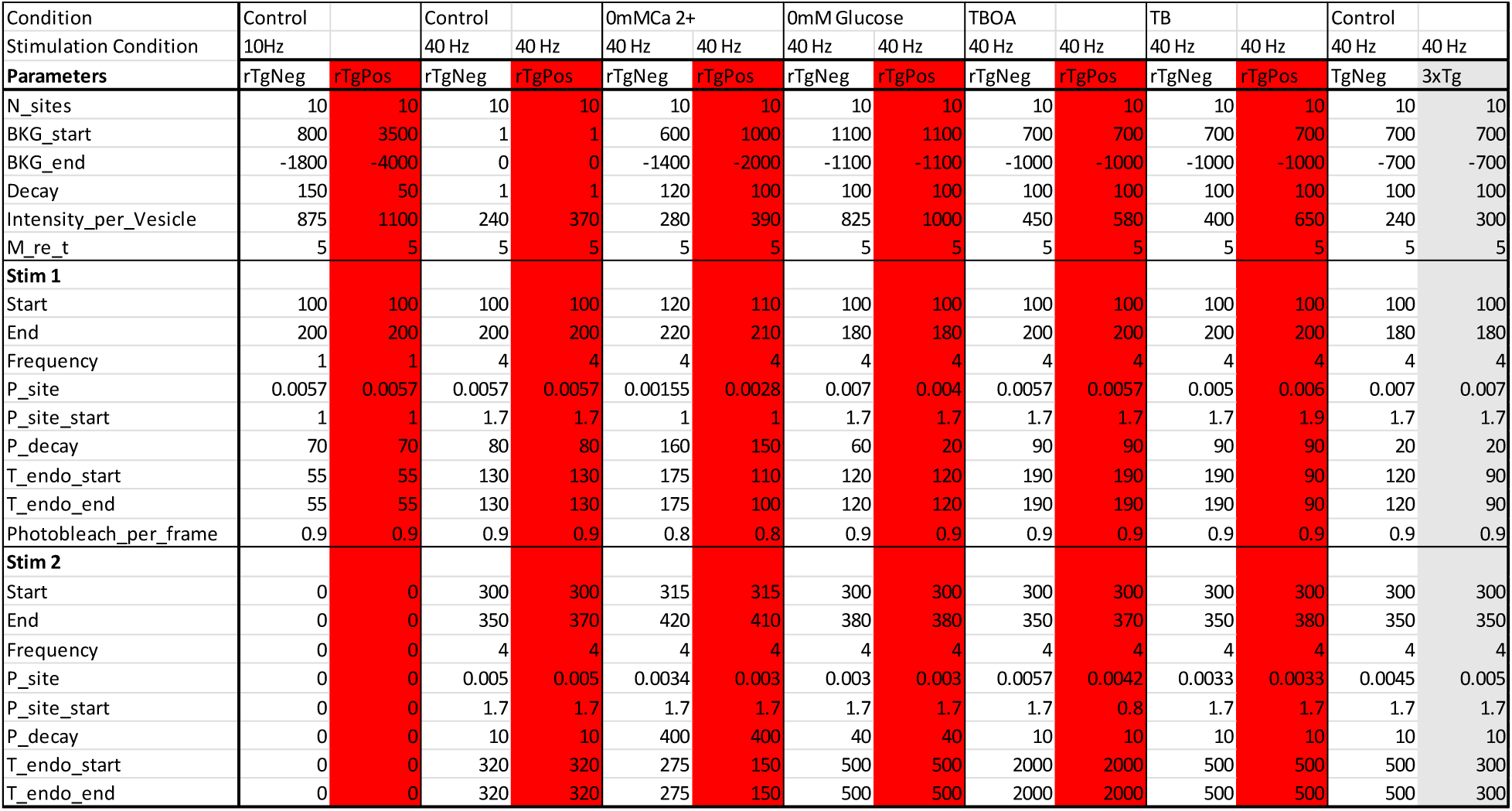
Computational Parameters used to model VGlut1-pHluorin stimulated intensity data.

## References

1. Hoover, B. R. et al. Tau Mislocalization to Dendritic Spines Mediates Synaptic Dysfunction Independently of Neurodegeneration. Neuron 68, 1067–1081 (2010).

2. Hunsberger, H. C., Rudy, C. C., Batten, S. R., Gerhardt, G. A. & Reed, M. N. P301L tau expression affects glutamate release and clearance in the hippocampal trisynaptic pathway. Journal of Neurochemistry 132, 169–182 (2014).

3. Kazim, S. F. et al. Early-Onset Network Hyperexcitability in Presymptomatic Alzheimer’s Disease Transgenic Mice Is Suppressed by Passive Immunization with Anti-Human APP/Aβ Antibody and by mGluR5 Blockade. Frontiers in Aging Neuroscience 9, (2017).

4. Ghatak, S. et al. Mechanisms of hyperexcitability in Alzheimer’s disease hiPSC-derived neurons and cerebral organoids vs isogenic controls. eLife 8, e50333 (2019).

5. Holth, J. K. et al. Tau Loss Attenuates Neuronal Network Hyperexcitability in Mouse and *Drosophila* Genetic Models of Epilepsy. J. Neurosci. 33, 1651 (2013).

6. McInnes, J. et al. Synaptogyrin-3 Mediates Presynaptic Dysfunction Induced by Tau. Neuron 97, 823–835.e8 (2018).

7. Zhou, L. et al. Tau association with synaptic vesicles causes presynaptic dysfunction. Nature Communications 8, 15295 (2017).

8. Taipala, E., Pfitzer, J. C., Hellums, M., Reed, M. N. & Gramlich, M. W. rTg(TauP301L)4510 mice exhibit increased VGlut1 in hippocampal presynaptic glutamatergic vesicles and increased extracellular glutamate release. Frontiers in Synaptic Neuroscience 14, (2022).

9. Siano, G. et al. Tau-dependent HDAC1 nuclear reduction is associated with altered VGluT1 expression. Frontiers in Cell and Developmental Biology 11, (2023).

10. Hunsberger, H. C. et al. Riluzole rescues glutamate alterations, cognitive deficits, and tau pathology associated with P301L tau expression. Journal of Neurochemistry 135, 381–394 (2015).

11. Pulido, C. & Ryan, T. A. Synaptic vesicle pools are a major hidden resting metabolic burden of nerve terminals. Science Advances 7, eabi9027.

12. Manus W. Ward, A. Cristina Rego, Bruno G. Frenguelli, & David G. Nicholls. Mitochondrial Membrane Potential and Glutamate Excitotoxicity in Cultured Cerebellar Granule Cells. J. Neurosci. 20, 7208 (2000).

13. Heine, K. B., Parry, H. A. & Hood, W. R. How does density of the inner mitochondrial membrane influence mitochondrial performance? *American Journal of Physiology-Regulatory*, Integrative and Comparative Physiology 324, R242–R248 (2023).

14. Kühlbrandt, W. Structure and function of mitochondrial membrane protein complexes. BMC Biology 13, 89 (2015).

15. Wolf, D. M. et al. Individual cristae within the same mitochondrion display different membrane potentials and are functionally independent. The EMBO Journal 38, e101056 (2019).

16. Gerencser, A. A. et al. Quantitative measurement of mitochondrial membrane potential in cultured cells: calcium-induced de- and hyperpolarization of neuronal mitochondria. The Journal of Physiology 590, 2845–2871 (2012).

17. Hu, Y. et al. Tau accumulation impairs mitophagy via increasing mitochondrial membrane potential and reducing mitochondrial Parkin. Oncotarget 7, 17356–17368 (2016).

18. Esteras, N. & Abramov, A. Y. Mitochondrial Calcium Deregulation in the Mechanism of Beta-Amyloid and Tau Pathology. Cells 9, (2020).

19. Keisuke Hirai et al. Mitochondrial Abnormalities in Alzheimer’s Disease. J. Neurosci. 21, 3017 (2001).

20. Chou, J. L. et al. Early dysregulation of the mitochondrial proteome in a mouse model of Alzheimer’s disease. Journal of Proteomics 74, 466–479 (2011).

21. Rhein, V. et al. Amyloid-β and tau synergistically impair the oxidative phosphorylation system in triple transgenic Alzheimer’s disease mice. Proceedings of the National Academy of Sciences 106, 20057–20062 (2009).

22. Trushina, N. I., Bakota, L., Mulkidjanian, A. Y. & Brandt, R. The Evolution of Tau Phosphorylation and Interactions. Frontiers in Aging Neuroscience 11, (2019).

23. DuBoff, B., Götz, J. & Feany, M. B. Tau Promotes Neurodegeneration via DRP1 Mislocalization In Vivo. Neuron 75, 618–632 (2012).

24. Li, X.-C. et al. Human wild-type full-length tau accumulation disrupts mitochondrial dynamics and the functions via increasing mitofusins. Scientific Reports 6, 24756 (2016).

25. Akbar, M. et al. Mitochondrial dysfunction and cell death in neurodegenerative diseases through nitroxidative stress. Brain Research 1637, 34–55 (2016).

26. Wang, W. et al. Damaged mitochondria coincide with presynaptic vesicle loss and abnormalities in alzheimer’s disease brain. Acta Neuropathologica Communications 11, 54 (2023).

27. Martínez-Reyes, I. et al. TCA Cycle and Mitochondrial Membrane Potential Are Necessary for Diverse Biological Functions. Molecular Cell 61, 199–209 (2016).

28. Lavie, J. et al. Ubiquitin-Dependent Degradation of Mitochondrial Proteins Regulates Energy Metabolism. Cell Reports 23, 2852–2863 (2018).

29. Ravanelli, S., den Brave, F. & Hoppe, T. Mitochondrial Quality Control Governed by Ubiquitin. Frontiers in Cell and Developmental Biology 8, (2020).

30. Jing, C. -h., et al. Autophagy activation is associated with neuroprotection against apoptosis via a mitochondrial pathway in a rat model of subarachnoid hemorrhage. Neuroscience 213, 144–153 (2012).

31. Liu, L. et al. Trans-Synaptic Spread of Tau Pathology In Vivo. PLOS ONE 7, e31302 (2012).

32. Maschi, D., Gramlich, M. W. & Klyachko, V. A. Myosin V functions as a vesicle tether at the plasma membrane to control neurotransmitter release in central synapses. eLife 7, e39440 (2018).

33. Voglmaier, S. M. et al. Distinct Endocytic Pathways Control the Rate and Extent of Synaptic Vesicle Protein Recycling. Neuron 51, 71–84 (2006).

34. Chanaday, N. L. & Kavalali, E. T. Optical detection of three modes of endocytosis at hippocampal synapses. eLife 7, e36097 (2018).

35. Reers, M., Smith, T. W. & Chen, L. B. J-aggregate formation of a carbocyanine as a quantitative fluorescent indicator of membrane potential. Biochemistry 30, 4480–4486 (1991).

36. Lanore, F. & Silver, R. A. Extracting quantal properties of transmission at central synapses. Neuromethods 113, 193– 211 (2016).

37. Reid, C. A. & Clements, J. D. Postsynaptic expression of long-term potentiation in the rat dentate gyrus demonstrated by variance-mean analysis. The Journal of Physiology 518, 121–130 (1999).

38. Scheuss, V. & Neher, E. Estimating Synaptic Parameters from Mean, Variance, and Covariance in Trains of Synaptic Responses. Biophysical Journal 81, 1970–1989 (2001).

39. Smith, H. L. et al. Mitochondrial support of persistent presynaptic vesicle mobilization with age-dependent synaptic growth after LTP. eLife 5, e15275 (2016).

40. Lopez-Manzaneda, M., Fuentes-Moliz, A. & Tabares, L. Presynaptic Mitochondria Communicate With Release Sites for Spatio-Temporal Regulation of Exocytosis at the Motor Nerve Terminal. Frontiers in Synaptic Neuroscience 14, (2022).

41. Joubert, F. & Puff, N. Mitochondrial Cristae Architecture and Functions: Lessons from Minimal Model Systems. Membranes 11, (2021).

42. Stephan, T., Roesch, A., Riedel, D. & Jakobs, S. Live-cell STED nanoscopy of mitochondrial cristae. Scientific Reports 9, 12419 (2019).

43. Wolf, S. G. et al. 3D visualization of mitochondrial solid-phase calcium stores in whole cells. eLife 6, e29929 (2017).

44. Sivandzade, F., Bhalerao, A. & Cucullo, L. Analysis of the Mitochondrial Membrane Potential Using the Cationic JC-1 Dye as a Sensitive Fluorescent Probe. Bio-protocol 9, e3128 (2019).

45. Leitz, J. & Kavalali, E. T. Ca^2+^ Influx Slows Single Synaptic Vesicle Endocytosis. J. Neurosci. 31, 16318 (2011).

46. Maschi, D. & Klyachko, V. A. Spatiotemporal Regulation of Synaptic Vesicle Fusion Sites in Central Synapses. Neuron 94, 65–73.e3 (2017).

47. Harris, J. J., Jolivet, R. & Attwell, D. Synaptic Energy Use and Supply. Neuron 75, 762–777 (2012).

48. Garcia, G. C. et al. Mitochondrial morphology provides a mechanism for energy buffering at synapses. Scientific Reports 9, 18306 (2019).

49. Alabi, A. A. & Tsien, R. W. Synaptic vesicle pools and dynamics. Cold Spring Harb Perspect Biol 4, a013680–a013680 (2012).

50. Crimins, J. L., Rocher, A. B. & Luebke, J. I. Electrophysiological changes precede morphological changes to frontal cortical pyramidal neurons in the rTg4510 mouse model of progressive tauopathy. Acta Neuropathologica 124, 777–795 (2012).

51. Klyachko, V. A. & Stevens, C. F. Excitatory and Feed-Forward Inhibitory Hippocampal Synapses Work Synergistically as an Adaptive Filter of Natural Spike Trains. PLOS Biology 4, e207 (2006).

52. Rolfe, D. F. S., Hulbert, A. J. & Brand, M. D. Characteristics of mitochondrial proton leak and control of oxidative phosphorylation in the major oxygen-consuming tissues of the rat. Biochimica et Biophysica Acta (BBA) - Bioenergetics 1188, 405–416 (1994).

53. Chinopoulos, C., Kiss, G., Kawamata, H. & Starkov, A. A. Chapter Seventeen - Measurement of ADP–ATP Exchange in Relation to Mitochondrial Transmembrane Potential and Oxygen Consumption. in Methods in Enzymolog*y* (eds. Galluzzi, L. & Kroemer, G.) vol. 542 333–348 (Academic Press, 2014).

54. Gramlich, M. W. & Klyachko, V. A. Nanoscale Organization of Vesicle Release at Central Synapses. Trends in Neurosciences 42, 425–437 (2019).

55. Vos, M., Lauwers, E. & Verstreken, P. Synaptic Mitochondria in Synaptic Transmission and Organization of Vesicle Pools in Health and Disease. Frontiers in Synaptic Neuroscience 2, (2010).

56. Pathak, D. et al. The role of mitochondrially derived ATP in synaptic vesicle recycling. J Biol Chem 290, 22325– 22336 (2015).

57. Díaz-García, C. M. et al. The distinct roles of calcium in rapid control of neuronal glycolysis and the tricarboxylic acid cycle. eLife 10, e64821 (2021).

58. Li, H. et al. Neurons require glucose uptake and glycolysis in vivo. Cell Reports 42, (2023).

59. Calvo-Rodriguez, M., Kharitonova, E. K. & Bacskai, B. J. In vivo brain imaging of mitochondrial Ca2+ in neurodegenerative diseases with multiphoton microscopy. Biochimica et Biophysica Acta (BBA) - Molecular Cell Research 1868, 118998 (2021).

60. Wilson, N. R. et al. Presynaptic Regulation of Quantal Size by the Vesicular Glutamate Transporter VGLUT1. J. Neurosci. 25, (2005).

61. Hu, C. et al. OPA1 and MICOS Regulate mitochondrial crista dynamics and formation. Cell Death & Disease 11, 940 (2020).

62. Schulz, K. L. et al. A New Link to Mitochondrial Impairment in Tauopathies. Molecular Neurobiology 46, 205–216 (2012).

63. Westermann, B. Mitochondrial fusion and fission in cell life and death. Nature Reviews Molecular Cell Biology 11, 872–884 (2010).

64. DeVos, S. L. et al. Synaptic Tau Seeding Precedes Tau Pathology in Human Alzheimer’s Disease Brain. Frontiers in Neuroscience 12, (2018).

65. Oddo, S. et al. Triple-Transgenic Model of Alzheimer’s Disease with Plaques and Tangles: Intracellular Aβ and Synaptic Dysfunction. Neuron 39, 409–421 (2003).

66. Shimamoto, K. et al. dl-*threo*-β-Benzyloxyaspartate, A Potent Blocker of Excitatory Amino Acid Transporters. Mol Pharmacol 53, 195 (1998).

67. Rose, C. R. et al. Astroglial Glutamate Signaling and Uptake in the Hippocampus. Frontiers in Molecular Neuroscience 10, (2018).

68. Srivastava, I., Vazquez-Juarez, E. & Lindskog, M. Reducing Glutamate Uptake in Rat Hippocampal Slices Enhances Astrocytic Membrane Depolarization While Down-Regulating CA3–CA1 Synaptic Response. Frontiers in Synaptic Neuroscience 12, (2020).

